# *In vivo* imaging of central nervous system regeneration using *Danionella cerebrum*

**DOI:** 10.1101/2025.05.13.653880

**Authors:** Mallika Misyuk, Joseph L. Austin, Noelle R. Walechka, Juliet A. Peterka, Matthew B. Veldman, Pui-Ying Lam

## Abstract

Rebuilding functional neuronal circuits after injury in the adult central nervous system (CNS) is unachievable for many vertebrates. In pro-regenerative models, it is unclear how regeneration and re-wiring is achieved in the CNS over long distances. The size and opacity of the adult vertebrate brain makes it difficult to study re-innervation patterns and dynamic cellular interactions during long-distance axon regeneration. Here, we harnessed the properties of the small and transparent adult *Danionella cerebrum* for longitudinal *in vivo* imaging of retinal ganglion cell axon regeneration, correlating cellular events with functional recovery. Our results suggest that, following optic nerve injury, the arborization pattern of re-innervation differs after regeneration, suggesting that new axon tracts are formed to restore functional vision. Additionally, myelin is not restored to pre-injury levels, even after functional recovery is achieved. The *D. cerebrum* model provides a unique opportunity to visualize and experimentally manipulate the spatial and temporal events during CNS regeneration in intact adult vertebrates.

**Research highlights:**

- Reveals the neuro-regenerative potential of the small, transparent adult vertebrate
- *Danionella cerebrum*, a novel model for central nervous system (CNS) repair studies.
- Demonstrates CNS regeneration using two visual functional recovery assays and longitudinal *in vivo* imaging of a novel adult axonogenesis reporter transgenic line, a myelin reporter line, and mosaic labeling of retinal ganglion cell axons.
- Establishes a method to compare retinal ganglion cell axons before and after CNS injury in adult *D. cerebrum*.
- Positions *D. cerebrum* as a powerful platform for visualizing long-distance CNS regeneration in intact adult vertebrates.

**Graphical abstract:** **Longitudinal live imaging of adult *D. cerebrum* reveals the formation of a new axon and myelin pattern post optic nerve injury along with functional recovery.**

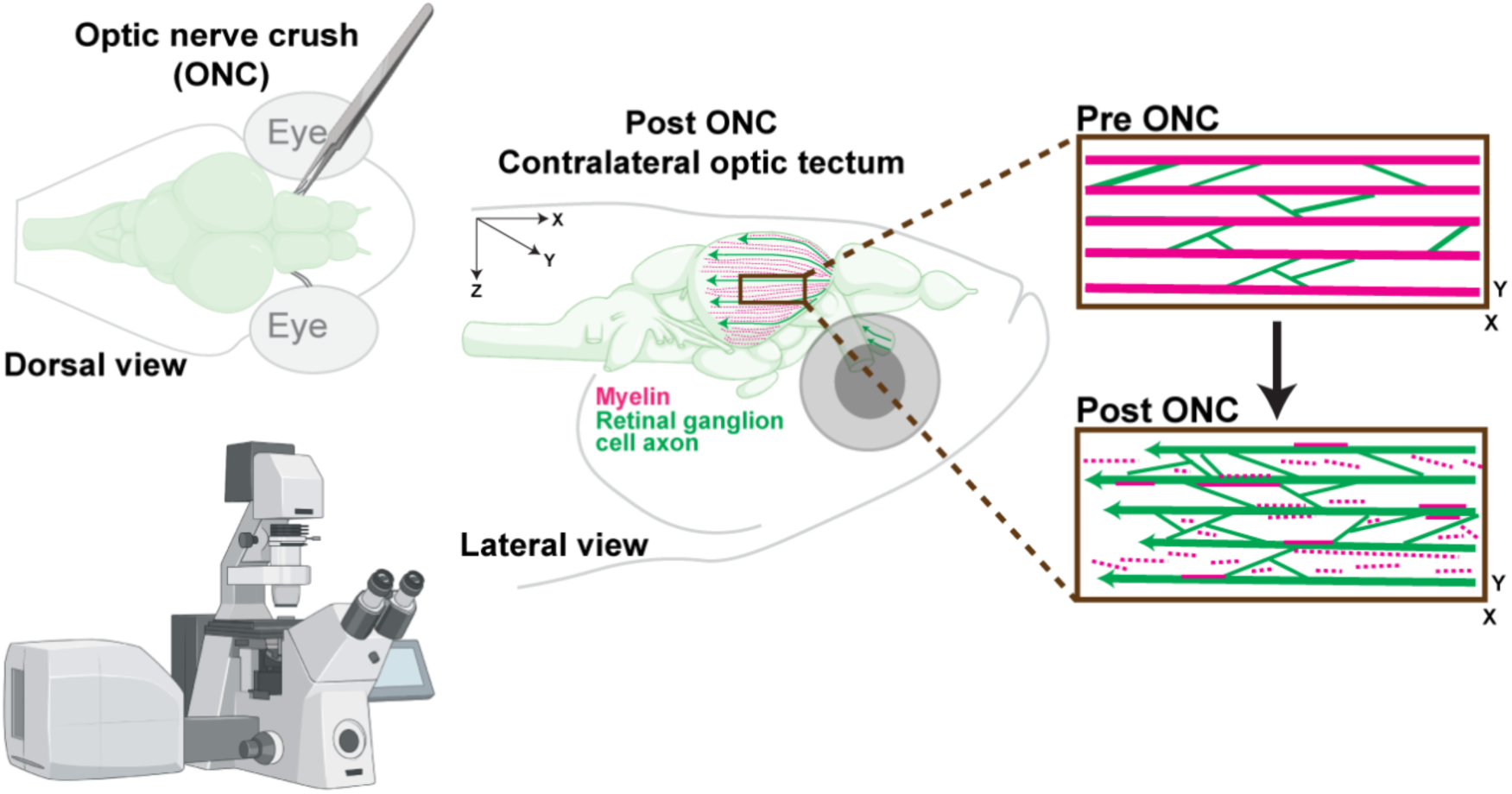

## Introduction

It is well known that after a central nervous system (CNS) injury, most mammals fail to sufficiently regenerate damaged axons to achieve proper target innervation. Long distance axon regeneration after an optic nerve injury has been successfully induced in mice using genetic manipulations (Park et al., 2008; Belin et al., 2015; Lim et al., 2016; Schaeffer et al., 2023), however, functional recovery of vision fails since axons do not make proper connections. Vision is restored after injury in pro-regenerative models, but it remains unclear how the neuronal arborization patterning is re-established during regeneration, nor is it known how axonal remodeling events during late regeneration contribute to functional recovery.

In nerve regeneration studies using vertebrate animals, the identification of key morphological events traditionally requires histological sectioning of fixed tissue, and/or tissue clearing (Chato-Astrain et al., 2022; Daeschler et al., 2022). Although optical tissue clearing techniques have enabled three-dimensional imaging of entire organs, it has not been possible to image dynamic cell-cell interactions, monitor the same animal over time, and correlate cellular events with functional recovery. Alternatively, the use of intra-vital microscopy in the CNS of traditional adult animal models requires the invasive implantation of an imaging window, yet allows for a limited imaging field and depth (Suthya et al., 2024). To overcome this, larval zebrafish have been used for the *in vivo* study of cellular dynamics and regeneration, owing to their small size and transparency. However, larvae lack complex memory-based behaviors and fully developed endocrine and neuro-immune systems, which makes it challenging to distinguish between different phases of regeneration and functional recovery or to identify sex-based differences and neuro-immune responses during regeneration (Murashova and Dyachuk, 2025). *Danionella cerebrum* (DC) is a new vertebrate model that provides a unique opportunity to visualize the entire brain, *in vivo,* at single cell resolution. Adult DC remain largely optically transparent, have the smallest known vertebrate brain, and lack a dorsal skull cap (Penalva-Tena et al.; Penalva et al., 2018; Schulze et al., 2018a; Britz et al., 2021; Lam, 2022; Lindemann et al., 2022; Rajan et al., 2022; Lee and Briggman, 2023; Cook et al., 2024; Groneberg et al., 2024; Vasconcelos et al., 2024; Veith et al., 2024; Zada et al., 2024), which makes DC particularly suitable for imaging cellular dynamics, especially in the CNS, in a live animal without the need for surgical manipulations.

Investigating dynamic cellular biology during regeneration remains challenging in traditional animal models due to limitations in size and transparency. The aim of this study was to establish DC as a new model for longitudinal *in vivo* investigation of CNS regeneration and functional recovery. We induced axonal injury in adult DC retinal ganglion cells (RGCs) by adapting the optic nerve crush (ONC) technique, which has been used in other animal models to study CNS regeneration (Bormann et al., 1998; Williams et al., 2020). To assess the regenerative capacity of adult DC, we performed longitudinal live imaging using novel cell-specific fluorescent reporter lines and correlated functional recovery with cellular events *in vivo*. Our findings demonstrate that the DC model offers new opportunities for future studies aimed at understanding how dynamic cellular events during regeneration contribute to functional recovery.

## Methods

### Fish husbandry and maintenance

All animal work was carried out in accordance with the National Institutes of Health Guide for Care and Use of Experimental Animals and approved by the Medical College of Wisconsin’s Institutional Animal Care and Use Committee protocol #AUA00007421 on 02/11/2023. All DC (obtained from a cross between DC at the Bolton Museum Aquarium, Bolton, UK and wild-caught samples purchased from The Wet Spot Tropical Fish store at Portland, OR, USA) were maintained at 28°C on a 14-10 hour light-dark cycle. DC larvae were co-cultured with euryhaline rotifers (*Brachonius plicatilis*) between 5 and 11 days post-fertilization (dpf). Thereafter, they were placed into a re-circulating water system and fed rotifers once a day and *Artemia nauplii* thrice a day until 14 dpf. Fish ≥ 1-month post-fertilization (mpf) were considered adults and were fed with freshly hatched *Artemia nauplii* twice a day. Spawning was encouraged by placing a ∼6 cm white opaque silicone tube (6 mm inner diameter × 8 mm outer diameter, McMaster-Carr, Elmhurst, IL, USA, Cat# 5054K812) on the bottom of tanks. After collection of embryos, the media in the plate was exchanged every 24 h with E3 containing 10 mM HEPES (Gold Biotechnology, Olivette, MO, USA, Cat# H-400-250).

### Generation of expression plasmids and transgenic lines

Tol2-*gap43:mGreenLantern-CAAX* was made by performing a Multisite Gateway LR reaction to assemble the following elements using LR Clonase II Plus enzyme (ThermoFisher Scientific, Waltham, MA, USA, Cat# 12538120): *p5E-gap43, pME-mGreenLantern*, *p3E-CAAX(tH)* (Addgene, Watertown, MA, USA, Cat# 109539) (Hall et al., 2020), and *pDestTol2pA2* (Kwan et al., 2007). *p5E-gap43* was created by amplifying the *gap43* promoter from plasmid *pW1GFG43SA* (Udvadia, 2008) using a polymerase chain reaction (PCR) with the following primers: Forward (F): ggggacaactttgtatagaaaagttggtcgactgtgtggcttccg, and Reverse (R): ggggactgcttttttgtacaaacttgggttcaagttggttctatcctccct. The resulting PCR fragment containing 3624 base pairs of the promoter, the transcription start site, and the 5’UTR sequence stopping immediately before the initial ATG codon was integrated into the pDONR^TM^P4-P1R vector using a Gateway BP reaction. *pME-mGreenLantern-CAAX* was created by amplifying *mGreenLantern* using *LifeAct-mGreenLantern* (Addgene Cat# 164459) (Campbell et al., 2020) as a template and primers F: ggggacaagtttgtacaaaaaagcaggctccaccatggtgagcaagggcgaggagctg, and R: ggggaccactttgtacaagaaagctgggtgcttgtacagctcgtccatgtcatg. The PCR product was then cloned into *pDONR221* using a Gateway BP reaction. The *Tol2-mbpa:TdTomato-CAAX* and *Tol2-isl2b:Tdtomato-CAAX* expression constructs were created using Gateway LR reactions and combining either *p5E-mbpa* or *p5E-isl2b* (Ben Fredj et al., 2010) with *pME-TdTomato-CAAX* (Addgene, Cat# 135204) (Oehlers et al., 2015), *p3E-polyA* (Tol2kit # 302) (Kwan et al., 2007) and *pDestTol2pA2* (Kwan et al., 2007). The zebrafish *mbpa* promoter (Jung et al., 2010; Pinzon-Olejua et al., 2017) was amplified from genomic DNA using primers F: ggggacaactttgtatagaaaagttggcttacactcactgacaaatag and R: ggggactgcttttttgtacaaacttgtctaggtgttgatctgttcag, then integrated into the *pDONR^TM^P4-P1R* vector using a Gateway BP reaction to create *p5E-mbpa*.

All single transgenic lines and sparsely labeled fish were generated by microinjection of 2-3 nL of a DNA-transposase mixture into 1-4 cell stage *wild-type* (*WT*) DC embryos. The DNA-transposase mixture contained 8-8.8 ng/µL of DNA, 12.6 ng/µL *in vitro* transcribed Tol2 transposase mRNA (mMESSAGE mMACHINE™ SP6 Transcription Kit, ThermoFisher Scientific, Cat# AM1340) and 0.05% phenol red (Sigma Aldrich, Burlington, MA, USA, Cat# P0290) as an indicator. Injected larvae were screened at 4 dpf and animals positive for reporter expression were raised to adults. To generate stable transgenic lines, adult F0 fish were in-crossed to produce F1 embryos that were screened for correct and complete transgene expression. Double transgenic lines were generated by outcrossing fish of the opposite sex from each transgenic line.

### Generation of microphthalmia-associated transcription factor (mitf) mutant line

Target sites for CRISPR/Cas9-directed mutagenesis were selected with the help of the online tool CHOPCHOP (Montague et al., 2014; Labun et al., 2016; Labun et al., 2019). A *mitf* gRNA (5’- GGGAGCCAAACTCAGCCCTC -3’) targeting exon 5 of the DC *mitf* gene was used to generate the *mitf* knockout. The generation of the sgRNA template and its subsequent transcription followed previously published protocols (Parvez *et al.,* 2021). Briefly, the DNA template for *in vitro* transcription was generated using a fill-in reaction of two annealed oligos. One oligo contained the *mitf* targeting sequence along with the SP6 promoter sequence (underlined) <u>ATTTAGGTGACACTATAG</u>GGAGCCAAACTCAGCCCTCGTTTTAGAGCTAGAAAT AGCAAG. The second oligo contained the constant region AAAAGCACCGACTCGGTGCCACTTTTTCAAGTTGATAACGGACTAGCCTTATTTTAACTT GCTATTTCTAGCTCTAAAAC. The fill-in reaction used the following parameters: 98°C – 2 min, 50°C - 10 min, 72°C - 10 min, 4°C - Hold. The annealed and filled-in oligo was purified using a Zymo DNA Clean and Concentrator-5 kit (Zymo Research, Irvine, CA, USA, Cat# D4013). *In vitro* transcription was performed using MEGAscript™ SP6 Transcription kit (ThermoFisher Scientific, Cat# AM1330) according to manufacturer’s guidelines. Samples were then purified using an RNA Clean and Concentrator-5 (Zymo Research, Cat# R1013). A 5 µL injection mixture was prepared with Buffer 3.1, 1000 ng gRNA, 0.84 µL EnGen Cas9 (NEB, Ipswich, MA, USA, Cat# M0646T). Then, 0.5 nL of the injection mixture was injected into 1-2 cell stage embryos. Genotyping of crispants, *WT* controls, and putative *mitf^−/-^*F1 animals was carried out by first extracting genomic DNA from tail fin clips (for adults) or from cells isolated by using a Zebrafish Embryo Genotyper (Lambert et al., 2018) (for larvae). PCR primers CCCACATGCCTAAACTTTGCAC and GTTGAGGGTGAGAAGGGCCA flanking the gRNA targeted site were used for PCR amplification of genomic DNA using Phire tissue direct PCR master mix (ThermoFisher Scientific, Cat# F170S). PCR products were then purified and sequenced.

### ONC and Optic Nerve Transection (ONT)

Adult DC were anaesthetized using 52.24 µg/mL Tricaine-S (ThermoFisher Scientific, Cat# NC0342409) diluted in fish system water. The fish was positioned on a custom 3-D printed (Lam 2022) holder at an oblique angle with the eye to be injured facing up. Excess fish water was removed. Under a dissecting microscope with bottom illumination, the optic nerve was exposed by gently removing the connective tissue on the dorsal half of the eye and rotating the eye out of its socket. Using a #5 forceps (Electron Microscopy Sciences, Hatfield, PA, USA, Cat# 0203-5-PS-1), ONC was carried out by closing the forceps on the optic nerve gently for 5 s while avoiding the ophthalmic artery posterior to the optic nerve. Similarly, ONT was carried out by cutting the optic nerve using micro-scissors (Fine Science Tools, Foster City, CA, USA, Cat# 15000-08). Immediately after the procedure, the ONC/ONT site was assessed under a dissecting microscope. A successful ONC was determined if the crushed site became opaque and the axonal tissue remained attached. A successful ONT was determined if the optic nerve head was fully detached from the rest of the optic nerve. The fish were then placed in a recovery tank for approximately 15 minutes or until the fish resumed normal swimming, after which they were transferred to the circulating tank system in the fish facility.

### Looming Response (LR) assay

To analyze the recovery of vision in adult DC post optic nerve injury, we adapted an LR assay, that was previously established in zebrafish to assess looming visual stimulus-evoked escape responses (Temizer et al., 2015; McKee and McHenry, 2020). The LR assay was performed in a custom-made blackout box measuring 45 cm × 38 cm × 30 cm. The box had a hinged front door and was lit with a uniform white LED light source (NXENTC A4 Tracing Light Pad, Amazon, Seattle, WA, USA, ASIN: B07D57F8X7) set to maximum brightness (1630 lx) to provide contrast for recording and tracking of fish. The fish was placed in a clear plastic tank (Michaels, Irving, TX, USA, Cat# 10728059) measuring 10.16 cm × 20.32 cm × 7.62 cm and filled with 770 mL of fish water maintained at 24 - 28°C. The bottom and lateral insides of the tank were covered in matte-clear tape (Gaffer Power Transparent Duct Tape, Amazon, ASIN: B076J2BS1B) to reduce internal reflection. The tank was placed in the blackout box 2.9 cm away from the LED light screen. The camera (Logitech, Lausanne, Switzerland, P/N: 860-000451, 1080p, 30fps) was placed 33 cm from the light screen to capture images of the whole tank. A Raspberry Pi monitor, 15×23.5 cm (ROADOM Raspberry Pi Screen, Amazon, ASIN: B09XDK2FRR), mirrored a computer monitor that played the stimulus video. The monitor was centered and placed face down at 2.54 cm above the water. The stimulus video was created using Adobe Premiere Pro 2024 (Adobe, San Jose, CA, USA, Version 24.0.1). The video projected a white screen for 2 min, followed by a looming stimulus presented as an expanding circle. The opacity, expansion speed and final size of the circle were optimized to illicit a robust change in swimming speed of uninjured *WT* fish while the stimulus appeared. An expanding opaque black circle that grew from 0 to 10.8 cm at a radius growth rate of 5.4 cm/s was used. The opaque black circle expanded over 2 s and remained on the screen for one minute. As the stimulus appeared, a grey rectangle appeared on the right side of the screen to allow for accurate tracking of the stimulus during analysis. This grey rectangle was reflected to the camera by a mirror (Iconikal Metal Compact, Amazon, ASIN: B0B75R1K8N) placed beside the tank.

Fish can respond to a looming stimulus even with one functional eye. To accurately assess the time-course of vision recovery post ONC, an ONT was first performed on the left eye of the fish to induce permanent blindness in that eye, followed by an ONC on the right eye of the same fish one day later. A LR assay was performed on the same fish over the course of 12 days to assess the recovery of vision on the ONC-injured eye. The fish was given adequate time (5-20 min) to acclimate to the looming tank environment before the looming assay was performed. Acclimation was considered complete when the fish displayed leisurely swimming throughout tank.

An ANY-Maze video tracking system (ANY-Maze, Dublin, Ireland, Version 7.4) was used for animal tracking. A Python code was created to analyze the data generated from ANY-Maze. The code identified the exact time the visual stimulus appeared by detecting the change in pixel intensity in the reflection of the mirror. The code averaged the swim speed of the fish 1 s before the stimulus and 1 second as the stimulus was played. 2-3 trials were conducted per fish for each time-point. Fish showed minimal looming responses at 1-day post ONC (dpONC). The average change in speed at this time-point was used as the baseline for determining a reaction score. If a trial’s average change in speed was greater than one standard deviation of the average change in speed at 1 dpONC, the trial scored a 1 and considered a positive response, otherwise, the trial scored a 0 and was considered a non-response. The score was averaged for the trials at each time-point for each fish. The scores were converted to a percentage and graphed using GraphPad Prism (GraphPad Software, San Diego, CA, USA, Version 10.4.1).

### Dorsal light response (DLR) assay

The DLR assay (Watanabe et al., 1989; Lindsey and Powers, 2007; Mensinger and Powers, 2007; Beckers et al., 2023) was adapted for DC and used to investigate the behavioral response of fish to visual stimuli by quantifying the tilt angle of the head orientation while swimming. Single fish were acclimated for 5 to 10 minutes (until normal swim behavior was observed) in a clear acrylic chamber (6 cm× 6 cm× 6 cm) containing 130 mL of fish water maintained at 24-26°C. Each apparatus had two side-by-side chambers, allowing for simultaneous recording of two fish. The lateral and bottom insides of the chamber were lined with 100% light blocking white vinyl (VELIMAX White Blackout Window Film, Amazon, ASIN: B08P8KQCL4), while the front was covered with one-way mirror film (KESPEN Window Privacy Film One Way, Amazon, ASIN: B07PBTWCXC) with 28% visible light transmission, oriented such that the reflective side faced inward. The back side of the chamber remained transparent. A uniform white LED light source (Invitrogen, Carlsbad, CA, USA, Cat# LB0100) illuminated the chamber from the back, operating at its maximum light intensity (17900 lx) setting to provide optimal contrast for video capture.

Fish were filmed swimming toward the mirror film using a camera (iPhone 11, 1080p resolution, 30 fps) positioned 12.5 cm away from the front of the chamber. The camera was focused on the center of the chamber, with fish monitored during recording to ensure head-on views were captured. Video recordings varied in length from less than 1 minute to approximately 5 minutes. For analysis, 3-5 frames were captured from each video recording, where the fish was positioned head-on relative to the camera. Head tilt angles were measured from the captured frames using Fiji (ImageJ Version 1.54p, NIH, Bethesda, MD, USA) (Schindelin et al., 2012). Specifically, the straight-line tool was used to draw a line connecting the top pixel of one eye to the top pixel of the other eye. This line was aligned at the intersection of a cross to facilitate the angle measurement. The angle was quantified using the angle tool in Fiji. The change in head tilt angle was normalized by subtracting the head tilt angle at pre ONC time-point to each time-point. Fish possessing a change in head tilt angle above 10 degrees at 1 dpONC were considered for further analysis. The change in head tilt angle was measured over several days and plotted using GraphPad Prism.

### Fish mounting and intubation for imaging

*For larval DC*: At the desired developmental stage, larval DC were anaesthetized using 200 µg/mL Tricaine-S diluted in E3 containing 10 mM HEPES. Anaesthetized larvae were mounted for imaging using 1% low melt agarose (Apex Bioresearch, Houston, TX, USA, Cat# 20-104) in E3 in the desired orientation on a 3.5 cm glass bottom petri dish (Cellvis, Mountain View , CA, USA, Cat# D35-20-0-N). Once the agarose had hardened, 2 mL of 200 µg/mL Tricaine-S in E3 was added to the petri dish to cover the agarose.

*For adult DC*: Fish were lightly anaesthetized with 65 µg/mL Tricaine-S diluted in fish water and then embedded in 4% low melt agarose dissolved in E3 buffered with 10mM HEPES on a custom imaging chamber equipped with an intubation inflow tube that was placed inside the mouth of the fish, supplying fresh fish water for the duration of imaging, as previously described (Lam, 2022). For long-term time-lapse imaging of regeneration, fish were mounted and then intubated with 65 µg/mL Tricaine-S diluted in fish water for the duration of imaging.

### Image acquisition

All images were acquired using the NIS elements software (Nikon, Minato City, Tokyo, Japan, Version 5.41.01) on a Nikon Ti2 inverted microscope (Nikon) fitted with a Yokogawa CSU-W1 spinning disk confocal head (Yokogawa Co., Hachioji City, Tokyo, Japan). Images were captured using a Prime 95B Scientific CMOS camera (Teledyne Vision Solutions, Waterloo, Ontario, Canada), and fluorophores were excited using a 488 nm or 561 nm excitation laser. Images were captured using the following objectives: 4× Air 0.2 Numerical aperture (N.A.), 10 × Air 0.45 N.A., 20× Air 0.75 N.A. or 40× Water 1.25 N.A.

### Analysis of gap43 reporter signal

All quantification was performed using Fiji (Schindelin et al., 2012) and graphs were generated using GraphPad Prism. For the analysis of developmental downregulation of *gap43* reporter signal in **Figure S4**, *gap43* reporter signal was quantified using maximum intensity projection (MIP) images from confocal z-stack images. A fixed region of interest (ROI) was defined in the hindbrain (HB), midbrain (MB), and forebrain (FB) using the bright-field image as guidance. Mean signal intensity in the ROI was calculated and normalized to 2 mpf using the formula: mean intensity in the ROI at X mpf time-point /2 mpf time-point. For the analysis of *gap43* reporter signal in response to an optic nerve injury in **Figure 3K, 3Q** and **Figure S5K, S5L**, *gap43* reporter signal was quantified using single z-plane images at the medial contralateral optic tectum (cTeO; ROI as shown by the white dotted line in **Figure 3B’-G’, B”-G”** and yellow dotted line in **O-P**). The mean intensity under the line-ROI indicated in **Figure 3J** was calculated at different time-points and normalized with pre ONC using the formula: mean intensity the under the line-ROI at X dpONC time-point/ pre ONC. For the analysis of *gap43* reporter signal in a z-plane where re-innervation was observed in the medial optic tectum (TeO) in **Figure 3L**, the intensity of *gap43* reporter signal under the line-ROI from the rostral to caudal cTeO was plotted.

### Analysis of oligodendrocyte membrane reporter signal

Quantification of oligodendrocyte membrane reporter signal was performed using Fiji (Schindelin et al., 2012) and graphs were generated using GraphPad Prism. The pattern of the oligodendrocyte membrane reporter signal shown in **Figure 6G** was analyzed by plotting the mean signal intensity under a line ROI in the MIP image of the dorsal cTeO. Quantification of the percentage area covered by the myelin signal in **Figure 6H** was performed in a defined ROI in the rostral-dorsal TeO using MIP images. The ROI was selected to ensure that no pigment spots were included in the quantification. The same signal thresholding parameters were applied for each animal to quantify the percentage of area covered by the oligodendrocyte membrane reporter *mbpa:TdTomato-CAAX*.

### Analysis of distribution of sparsely labeled RGCs pre- and post-injury by dept

Confocal z-stack images of sparsely labelled RGCs at pre- and post-ONC time points were quantified for signal intensity by depth using Fiji (Schindelin et al., 2012) and graphs were generated using GraphPad Prism. The post ONC image was registered to the pre ONC image using the “BigWarp” Fiji plug-in (Schindelin et al., 2012) (Thuma et al., 2023). A fixed ROI in the cTeO was used to plot the signal intensity by depth using the “plot z-axis profile” option in Fiji (Schindelin et al., 2012).

### Post-acquisition image processing and video rendering

Post-acquisition, images in **Figure 3H-I**, and **Figs. 4-6** were subjected to Denoise.ai (NIS elements, Version 5.41.01). For image registration across different time-points, an image from the brightfield channel was used as a reference. Images in **Figs. 3-6** were registered in the X-Y direction using the “Align Image by Line ROI” Fiji plug-in. Additionally, in **Figure 4B-E, 4B’-E’** and **5E-N**, the “BigWarp” Fiji plug-in was also used for image alignment. In **Supplemental Video 4**, video alignment was done using the FiJi plug-in “Linear Stack Alignment with SIFT”(Lowe, 2004). In **Supplemental Video 5** and **6**, Denoise.ai and 3-D deconvolution (NIS elements; parameters used were: “Deconvolution Method”: Automatic, “Modality”: Spinning Disk, “Pinhole Size”: 50.00 µm, “Magnification”: 40.0x, “Numerical Aperture”: 1.15, “Immersion Ref. Index”: 1.333, “Calibration”: 0.275 µm/px, “Z-step”: 0.400 µm) were applied. 3-D rendering videos were created using the “Movie Maker” function in NIS elements. Adobe Premiere Pro was used to add titles, annotations, timestamps, and to sync videos.

### Statistical analysis

Preliminary n number estimations were done using G*power (Kang, 2021) (Version 3.1.9.4; α = 0.05, and power (1-β) = 0.95). All statistical analysis post data acquisition was performed using GraphPad Prism. All data were tested for normal distribution using the Shapiro-Wilk normality test. 1-way Analysis of Variance (ANOVA) was carried out when comparing means of three or more independent groups. A Repeated Measures 1-way ANOVA was carried out when comparing means of three or more groups that are dependent. A Repeated Measures 2-way ANOVA was carried out when means of two or more independent groups were compared over time. The mixed effects ANOVA model was fitted when the sample size of each group was different.

## Results

### Adult D. cerebrum can functionally recover from an optic nerve injury

In non-mammalian vertebrates, RGC axons exit the eye as the optic nerve and innervate the TeO, the largest brain center for visual processing. After an optic nerve injury in zebrafish, approximately 75% of RGCs survive (Zou et al., 2013) with axonal regeneration (Udvadia, 2008; Diekmann et al., 2015) leading to functional recovery (Mensinger and Powers, 2007; Beckers et al., 2023; Hu and Veldman, 2025). It is not known if the CNS of adult DC can regenerate. We established the ONC technique in adult DC to inflict a CNS injury (**Figure 1A-A’**), then assessed their ability to regain vision using two behavioral assays, a DLR and a LR assay. The DLR assay has been used to assess vision in fish. When vision is lost in one eye, fish tend to tilt their body such that the eye that has lost vision faces upwards. The recovery of head tilt correlates with the recovery of vision (Watanabe et al., 1989; Lindsey and Powers, 2007; Mensinger and Powers, 2007; Beckers et al., 2023; Hu and Veldman, 2025). Based on this principle, we devised a DLR assay for adult DC to accommodate their small size and transparency for video recording and analysis (**Figure 1B, S1A and Methods**). After subjecting *WT* animals to an ONC, we measured the head tilt angle over the course of 18 days (**Figure 1B’ and 1C**). Compared to pre ONC, a significant change in head tilt angle was observed at 1 dpONC (**Figure 1D**; Total pre ONC vs 1 dpONC *P value* < 0.0001). Head tilt angle recovered by 18 dpONC, suggesting that adult DC can recover vision after an ONC (**Figure 1D and Supplemental Video 1**). When we compared the recovery of males to females, no significant difference was observed except at 10 dpONC (males vs females at 10 dpONC *P* = 0.0449).

**Figure 1.**
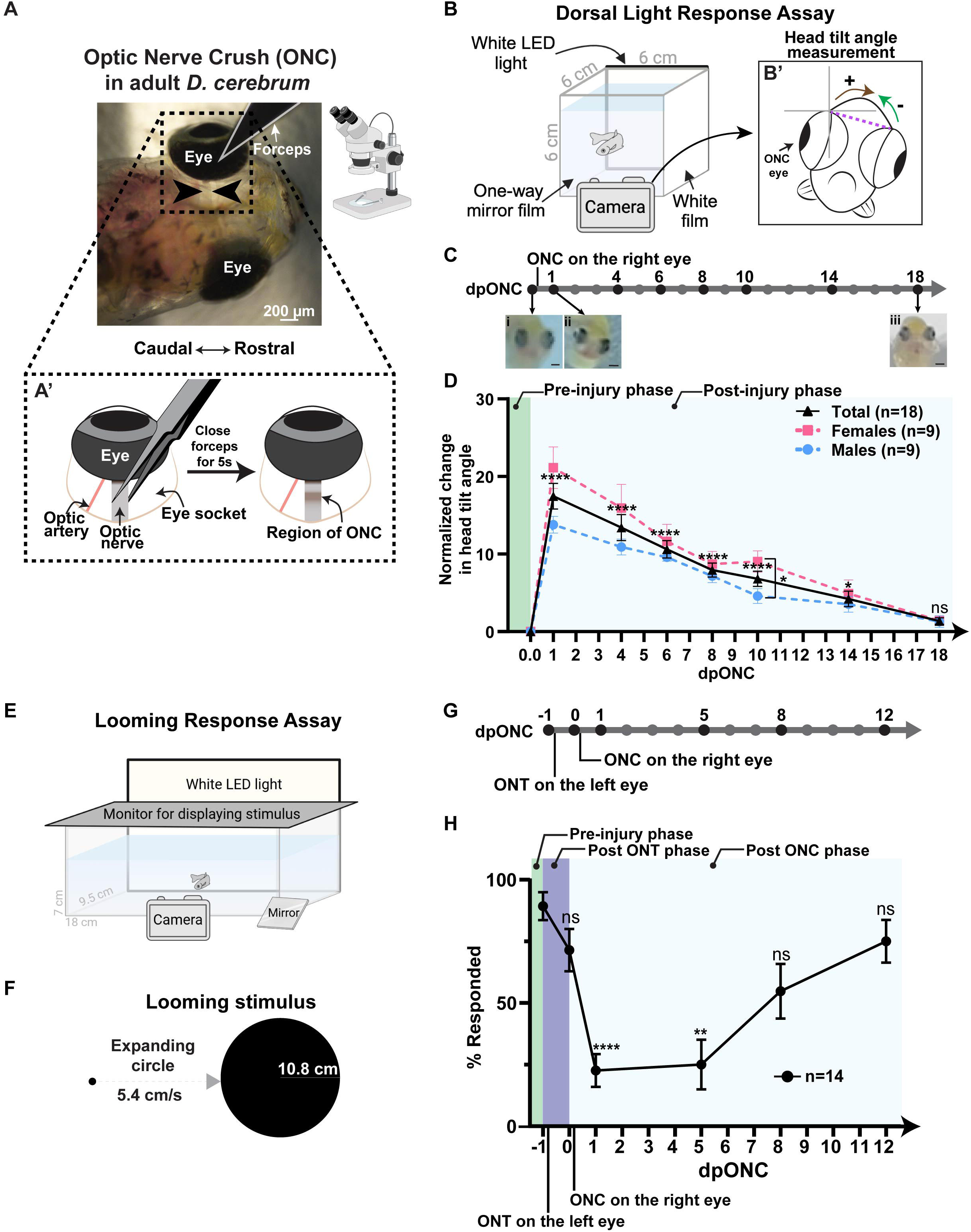
Assessing functional recovery after an ONC using DLR and LR Assays. **(A)** Dorsal view of an adult DC pre ONC with black arrowheads flanking the optic nerve. **(A’)** Schematic representation of the black dotted area in (A) illustrating the ONC procedure. After a successful ONC, the region of the optic nerve subjected to ONC turns opaque. **(B)** Schematic representation the DLR assay chamber. **(B’)** Illustration of how the head tilt angle is measured. Brown arrow showing a right tilt (positive angle) and green arrow showing a left tilt (negative angle). **(C)** Timeline schematic of the DLR assay with the time-points that head tilt angle was measured indicated by black dots. (i-iii) Representative images of the same fish (i) pre ONC, and at (ii) 1 dpONC and (iii) 18 dpONC. Scale bar = 200 µm. **(D)** Change in head tilt angle compared to pre ONC values. All data are represented in mean ± Standard Error of Mean (±SEM). Black solid line represents all animals (n = 18), pink dotted line represents females (n = 9), and blue dotted line represents males (n = 9). *P* values were calculated using Mixed Effects Repeated Measures 2-way ANOVA. *P* values of all animals (total) of each time-points are indicated above the trace where pre and post ONC measurements were compared. *P* values from left to right: *P* < 0.0001, *P* < 0.0001, *P* < 0.0001, *P* < 0.0001, *P* < 0.0001, *P* = 0.0105, and *P* = 0.1088. For females vs. males, at 1 dpONC, *P* = 0.0655; at 4 dpONC, *P* = 0.3086; at 6 dpONC, *P* = 0.6475; at 8 dpONC, *P* = 0.6981; at 10 dpONC *P* = 0.0449; at 14 dpONC *P* = 0.7762; and 18 dpONC *P* = 0.9985. Annotations as presented on graph: ns (not significant) *P* ≥ 0.05, * *P* < 0.05, ** *P* < 0.01, *** *P* < 0.001, **** *P* < 0.0001. **(E)** Schematic representation of the LR Assay. **(F)** Schematic illustration of the expanding circle parameters used in the LR assay. **(G)** Timeline schematic of the LR assay with the time-points that LR was assessed indicated by black dots. **(H)** Percentage of fish responding to the LR stimulus pre-and post-injury. All data represented in mean ±SEM (n = 14 including 7 females and 7 males). *P* value of each time-point was calculated using Repeated Measures 1-way ANOVA. *P* values represented as compared to -1 dpONC/pre-injury time-point. *P* values from left to right: *P* = 0.5013, *P* < 0.0001, *P* = 0.0017, *P* = 0.1126, and *P* = 0.7827. Annotations as presented on graph: ns *P* ≥ 0.05, ** *P* < 0.01, **** *P* < 0.0001.

To evaluate the functional vision of injured DC, we established an LR assay (**Figure 1E and F and S1B**). This assay tests the ability of fish to detect a predatory stimulus and has been used to assess visual function (Temizer et al., 2015; Dunn et al., 2016). Fish can display an escape behavior in response to a looming stimulus with only one eye. To specifically assess vision recovery from the ONC-injured eye, the optic nerve of one eye was first transected, then an ONC was performed on the other eye one day later (**Figure 1G**). We used the DLR assay to validate that vision did not recover after a full ONT in adult DC (**Figure S2;** Total pre ONT vs 18 dpONC *P value* < 0.0001). The percentage of fish responding to a looming stimulus at 1 dpONC was significantly lower compared to those that responded pre-injury (**Figure 1H**; -1 dpONC vs 1 dpONC *P value* < 0.0001). However, 22.62% of the fish were scored as responding to the stimulus at 1 dpONC despite having no intact optic nerves. This was likely due to sporadic movement of the fish coinciding with stimulus. Vision recovery was observed at 8 dpONC, although the percentage of fish that responded was still lower than at pre ONC levels (**Figure 1H and Supplemental Video 2**). The time to recover from ONC, as determined using the DLR assay differed from the LR assay. This might be due to the different synaptic connections required to elicit different visual responses as it was shown that teleost fish use different visual pathways when responding to different visual cues (Nevin et al., 2010). Nonetheless, our results show that adult DC can recover from an ONC injury, the progress of which can be assessed using a DLR and/or LR assay.

### An axonogenesis reporter line for adult fish

To facilitate the *in vivo* visualization of regenerating axons, we generated a novel transgenic DC line, *Tg(gap43:mGreenLantern-CAAX)*. Previous work has shown that the *gap43* promoter from Japanese pufferfish, *Takifugu,* can be used to track neuronal development and regeneration in zebrafish (Udvadia, 2008). At early developmental stages (2 dpf), robust *gap43* promoter-driven expression in the brain and eye was observed (**Figure 2A-D, 2A’-D’ and S3**) and persisted into the young adult stage (2.5 mpf) (**Figure 2E-E’**). *gap43* promoter-driven expression was detected in different regions of the FB, MB, and HB at all depths imaged, from the dorsal plane through to the ventral plane (**Figure 2G-I, 2G’-I’, Supplemental Video 3**). These observations demonstrated that this *gap43* promoter-based transgenic line shows expression throughout the brain and allows for monitoring of neurogenesis in the entire brain of adult DC.

**Figure 2.**
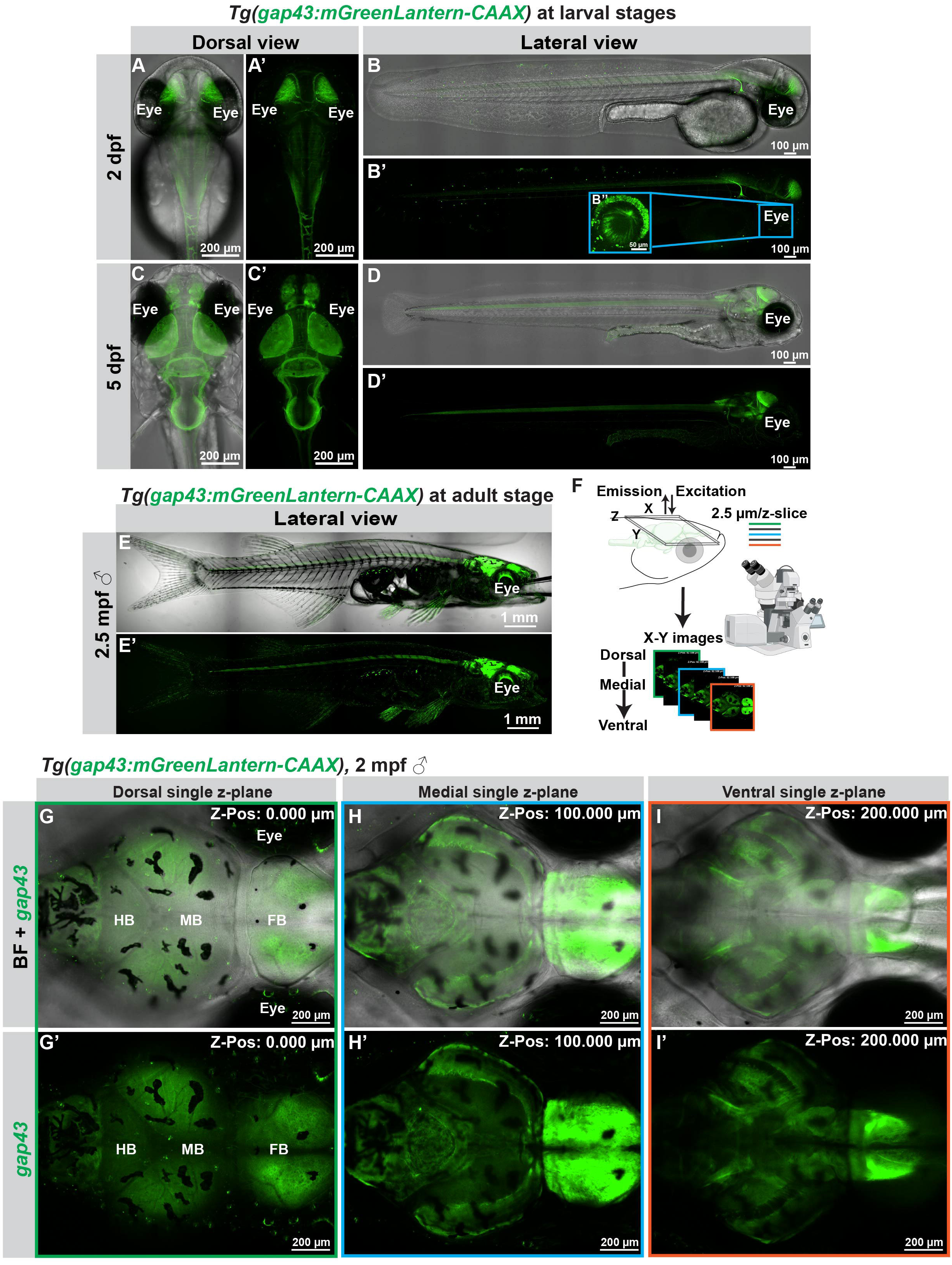
Generating the *Tg(gap43:mGreenLantern-CAAX)* axonogenesis reporter line. **(A-D)** and **(A’-D’)** Confocal images of *Tg(gap43:mGreenLantern-CAAX)* larvae acquired using a 20X objective (N.A. = 0.75, 0.9 µm/step). **(A-D)** *gap43* promoter driven expression (green channel) overlayed with the bright field (BF) channel in 2 and 5 dpf larvae. **(A-A’ and C-C’)** Dorsal confocal MIP images. The total z-stack was 199.80 µm in (A-A’) and 246.60 µm in (C-C’). (B-B’ and D-D’) Lateral confocal MIP images. The total z-stack was 199.90 µm in (B-B’) and 199.80 µm in (D-D’). (B’’) Inset of the region marked in the blue rectangle in B’ showing *gap43*-driven expression of fluorescent protein in the eye. **(E-E’)** Lateral confocal MIP images of an adult *Tg(gap43:mGreenLantern-CAAX)* male at 2.5 mpf. Image was acquired using a 4X objective (N.A. = 0.2). The total z-stack was 1050 µm with a 25 µm/step. ♂ = Male. **(E)** *gap43* promoter driven expression (green channel) overlayed with the BF channel. **(E’)** Green channel only. **(F)** Schematic representation of the approach used to acquire confocal images of the adult DC brain (images shown in G-I and G’-I’). Z-stacks were acquired at 2.5 µm/z-step using a 10X objective (N.A. = 0.45) on a 2 mpf *Tg(gap43:mGreenLantern-CAAX)* male measuring 1.32 cm in length. **(G-I)** *gap43* promoter driven expression (green channel) overlayed with the bright field (BF) channel. **(G’-I’)** Green channel only. **(G and G’)** Single plane image of the dorsal brain (z-position= 0.00 µm). **(H-H’)** Single plane image of the medial brain (z-position = 100.00 µm). **(I-I’)** Single plane image of the ventral brain (z-position = 200.00 µm).

**Figure 3.**
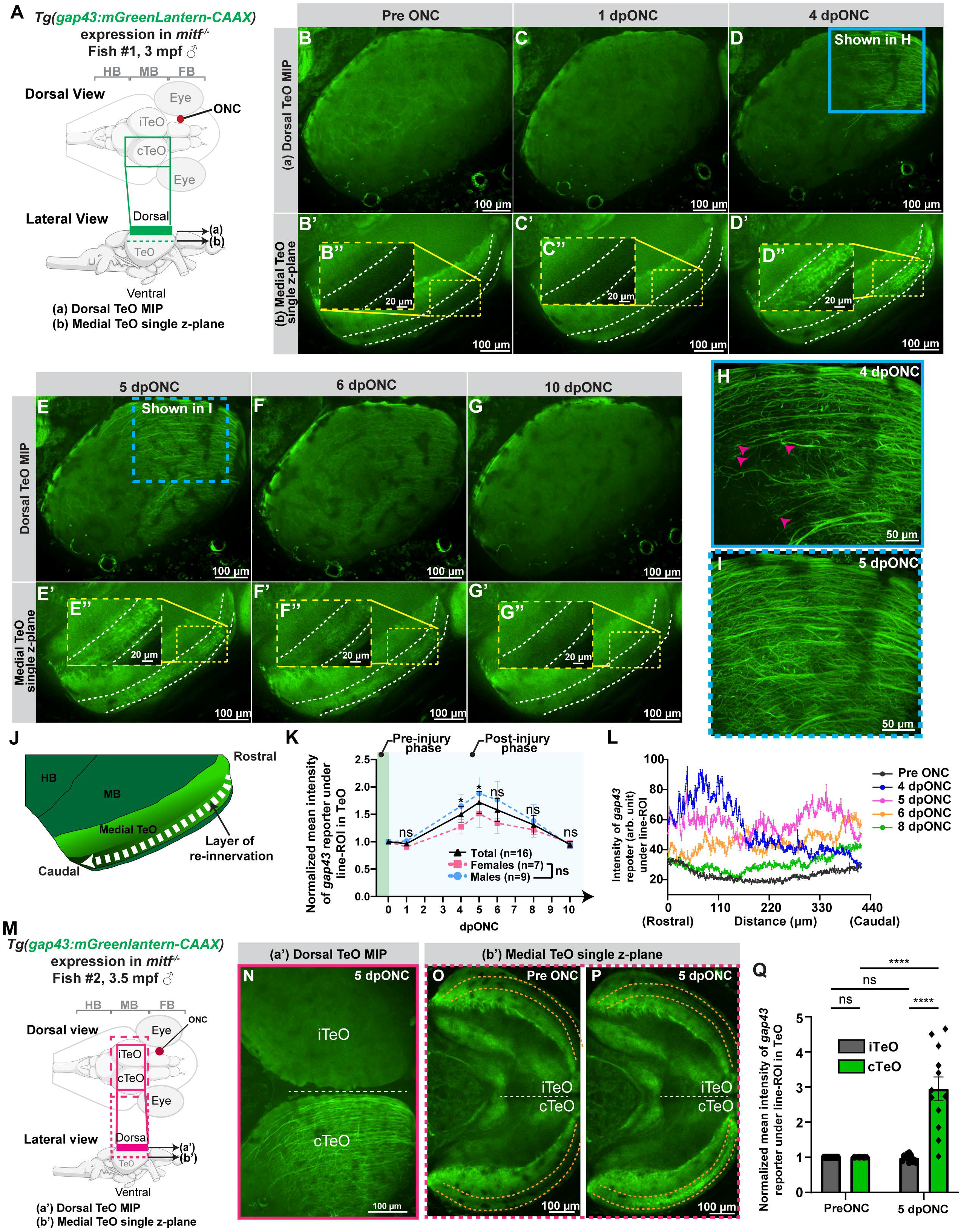
Longitudinal confocal imaging of axon regeneration after an ONC injury. **(A)** Schematic illustrations showing the imaged area depicted in B-G and B’-G’. The solid green rectangle in the lateral view (a), corresponds to the area imaged and presented as MIPs of the dorsal portion of the TeO in B-G. The green dotted line in the lateral view (b), corresponds to an area more ventral to (a) and presented as single plane images of the medial TeO in B’-G’. ♂ = Male. **(B-G, B’-G’, B”-G”)** Confocal images acquired in a *Tg(gap43:mGreenLantern-CAAX)* male *in mitf^−/-^* at 3 mpf with a length of 1.36 cm using a 20X objective (N.A. = 0.75, 0.9 µm/z-step). **(B-G)** Dorsal TeO MIP images of the *gap43* promoter-driven fluorescent signal at the indicated time-points. The total z-stack thickness was 117.00 µm in (B), 115.20 µm in (C), 119.70 µm in (D), 112.50 µm in (E), 115.20 µm in (F), and 118.80 µm in (G). **(B’-G’)** Single plane images of a deeper medial region in the TeO of the same fish. **(B”-G”)** Magnified area corresponding to the yellow dotted rectangle in B’-G’. Measurements of *gap43* reporter signal were made in the region bounded by the white dotted lines. **(H)** Magnified view of the region indicated by a blue rectangle in D. **(I)** Magnified view of the region indicated by a blue rectangle in E. **(J)** *gap43* reporter signal intensity was quantified in the area indicated by the white dotted line. This area represents the site of re-innervation (marked by black arrow). **(K)** Mean intensity of *gap43* reporter expression in the ROI indicated in (J) normalized to pre ONC levels was measured for 10 dpONC. All data represented in mean ±SEM. Black line represents all animals (n = 16), pink dotted line represents the females (n = 7), blue dotted line represents the males (n = 9). *P* values at each time-point compared to pre ONC were calculated using Mixed Effects Repeated Measures 2-way ANOVA. *P* values of all animals (total) from left to right: *P* = 0.9728, *P* = 0.0446, *P* = 0.0346, *P* = 0.1417, *P* = 0.5474, *P* = 0.9576. For females vs males, at 1 dpONC, *P* = 0.3399; at 4 dpONC, *P* = 0.3250; at 5 dpONC, *P* = 0.6274; at 6 dpONC, *P* = 0.5323; at 8 dpONC *P* = 0.8613; and at 10 dpONC *P* = 0.9375. Annotations as presented on graph: ns *P* ≥ 0.05, * *P* < 0.05. **(L)** Intensity of *gap43* reporter signal (arbitrary units) in the TeO ROI (indicated in J) measured in the same animal at the time-points indicated. **(M)** Schematic representation of regions and depths imaged in N-P. Magenta rectangle in the dorsal view shows the region imaged in (N) and the dotted rectangle shows the region in O-P. (a’) Solid bar corresponds to the region imaged and shown in (N) as a MIP. (b’) Dotted line corresponds to the single plane images shown in (O-P). **(N-P)** Confocal images acquired using a 10X objective (N.A. = 0.45, 2.5 µm/z-step) in a *mitf^−/-^Tg(gap43:mGreenLantern-CAAX)* male at 3.5 mpf with a length of 1.30 cm. White dotted line separates the cTeO and the iTeO. **(N)** Dorsal TeO MIP image of the expression of *gap43* reporter at 5 dpONC. The total z-stack thickness was 125.00 µm. **(O-P)** Medial TeO single z-plane image of the expression of *gap43* reporter at time-points indicated. The ROI quantified in Q is demarked by the orange dotted lines. **(Q)** Intensity of *gap43* reporter expression normalized to pre ONC levels in the ROI indicated in (O-P). All data represented in mean ±SEM (n = 12). *P* values calculated using Repeated Measures 2-way ANOVA. For pre ONC, iTeO vs cTeO *P* > 0.9999; for 5 dpONC, iTeO vs cTeO *P* < 0.0001; for iTeO, pre ONC vs 5 dpONC *P* = 0.9349; for cTeO; and pre ONC vs 5 dp ONC *P* < 0.0001. Annotations as presented on graph: ns *P* ≥ 0.05, **** *P* < 0.0001.

**Figure 4.**
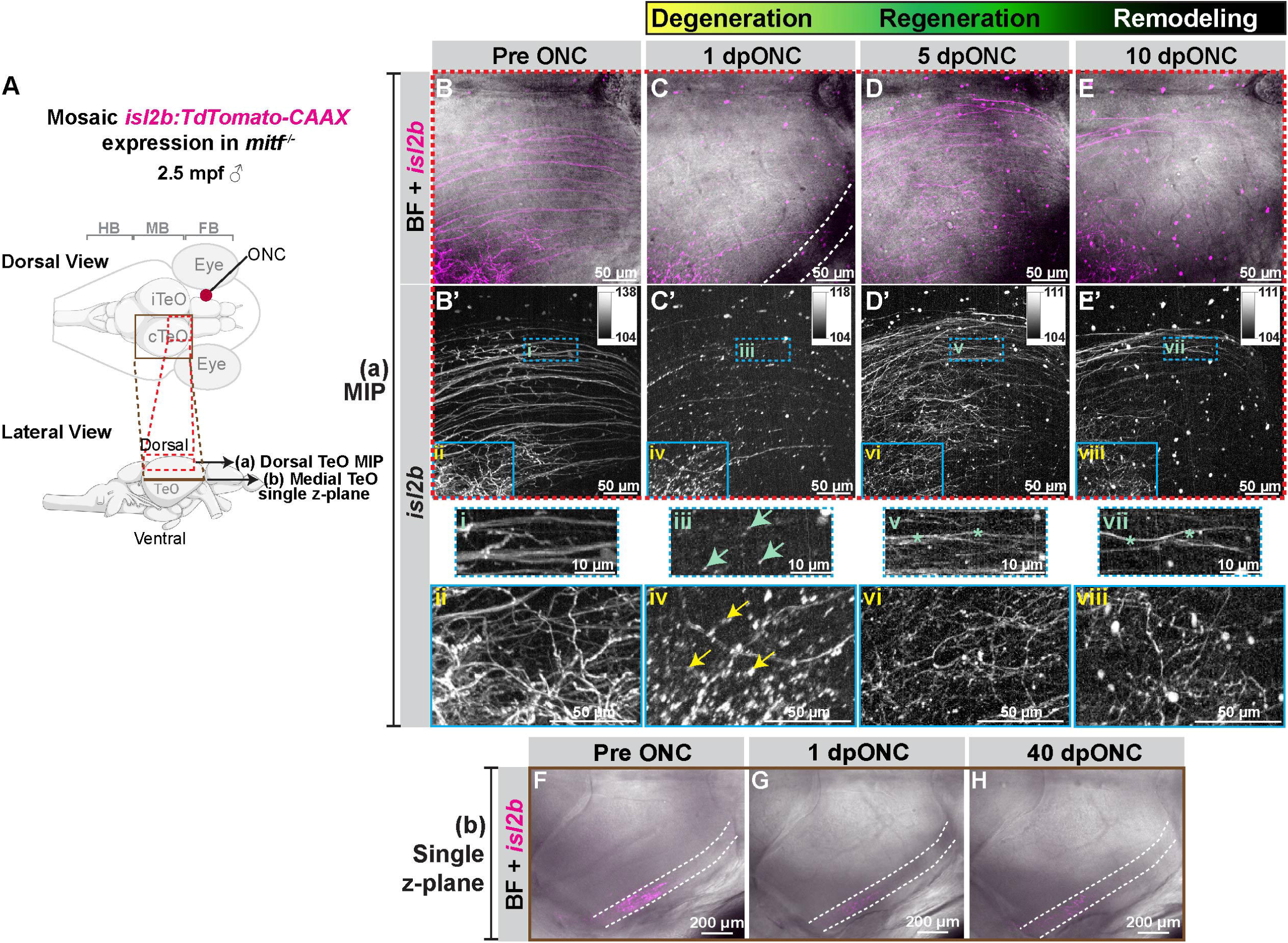
Assessing retinal ganglion cell axon degeneration and regeneration using a sparse labeling technique. **(A)** Schematic representation of regions imaged and shown in B-E and B’-E’. (a) Red dotted rectangle corresponds to the dorsal TeO area imaged and shown as a MIP in B-E and B’-E’. (b) Solid brown line indicates the position within the TeO imaged and shown as a single z-plane image in F-H. ♂ = Male. **(B-E)** and **(B’-E’)** All images were acquired using a 40X objective (N.A. = 1.15, 0.4 µm/z-step) in a male *mitf^−/-^* fish at 2.5 mpf 1.34 cm in length with mosaic *isl2b:TdTomato-CAAX* expression. The total z-stack thickness used for MIP was 112 µm. **(B-E)** MIP images showing the mosaic expression of *isl2b:TdTomato-CAAX* (magenta) combined with the corresponding bright field (BF) image in the dorsal region of the TeO at the time-points indicated. Dotted line in (C) indicates the location of the darker shade neuronal debris accumulation in the cTeO at 1 dpONC. **(B’-E’)** Grey scale images of B-E with look-up tables (LUTs) shown in the upper right inset. **(i-viii)** Magnified view of regions of interest as indicated by rectangles in B’-E’. Green and yellow arrows highlight degenerating *isl2b:TdTomato-CAAX* expressing neurites. Green asterisks highlight remodeling of *isl2b:TdTomato-CAAX* expressing neurites. **(F-H)** Single z-plane images of medial TeO region showing the mosaic expression of *isl2b:TdTomato-CAAX* (magenta) combined with the corresponding bright field (BF) image at the time-points indicated. Dotted line indicates the area of the TeO where *isl2b:TdTomato-CAAX* expressing neurons are innervating. Images were acquired using a 20X objective lens (N.A. = 0.75).

The expression of the endogenous *gap43* gene is downregulated as the animal ages due to the conclusion of neurogenesis (Reinhard et al., 1994; Karimi-Abdolrezaee et al., 2002). We performed longitudinal imaging experiments on the same *Tg(gap43:mGreenLantern-CAAX)* animals to quantify levels of *gap43* promoter-driven expression in multiple brain regions (**Figure S4A**). We observed significant downregulation of *gap43* reporter signal in all 3 regions of the brain at 6.5 mpf as compared to 2 mpf (for HB 2 mpf vs 6.5 mpf *P* value < 0.0001, for MB 2 mpf vs 6.5 mpf *P* value = 0.0012, for FB 2 mpf vs 6.5 mpf *P* value = 0.0007). Although levels had decreased in all regions tested, we could still detect signal at 6.5 mpf (**Figure S4B-J**), suggesting that fish at this age continue to exhibit some degree of neurogenesis.

### Axon regeneration in vivo

To visualize RGC axon regeneration *in vivo*, we performed an ONC in adult *Tg(gap43:mGreenLantern-CAAX)* fish then carried out longitudinal imaging on the same fish. We observed an upregulation of *gap43* reporter signal in the cTeO starting at 4 dpONC, followed by its downregulation by 9 dpONC (**Figure S5A-G**). Interestingly, when animals were subjected to an ONT, there was no significant difference in the *gap43* reporter signal intensity between pre ONT and 5 dpONT in the cTeO (**Figure S5H-K**). This suggests that a more severe transection injury does not lead to the appearance of regenerating neurites and corroborates our observation that animals receiving an ONT lack vision recovery when assessed using the DLR assay (**Figure S2**). We originally generated the *Tg(gap43:mGreenLantern-CAAX)* transgenic line in *WT* DC. The presence of melanophore pigment cells in the epidermal layer dorsal to the brain (**Figure S5D**), dynamically changed their shape and hindered our ability to quantify and compare the intensity of the *gap43* reporter signal in the same animal’s brain over time. To overcome this, we generated a DC pigmentation mutant for *in vivo* imaging of neuro-regeneration. Mutating the *mitfa* gene in zebrafish produced fish that lack melanophores (Lister et al., 1999). We identified the DC *mitf* gene using the NCBI DC Genome assembly ASM722483v1, then designed gRNAs to generate a *mitf^−/-^* mutant line using CRISPR-Cas9 mediated mutagenesis (**Figure S6A**). *mitf^−/-^*mutant DC have reduced melanophore pigment cells when compared to *WT* fish, both at larval (**Figure S6B and C)** and adult stages (**Figure S6D-I**). There was no significant difference in the looming response between *mitf^−/-^* and *WT* animals, suggesting that *mitf^−/-^* fish have normal vision (**Figure S6J**). We performed an ONC on *mitf^−/-^* animals and used the DLR assay to assess their ability to regenerate. Results between *WT* and *mitf^−/-^* animals were similar (**Figure S6K**). One difference we did observe, however, was that *mitf^−/-^* animals displayed a significantly slower swim speed in response to a looming stimulus when compared to *WT* animals (**Figure S6L**; post stimulus *WT* vs *mitf^−/-^, P* value *=* 0.0001).

Pigment-deficient *mitf^−/-^* fish were used to generate a *Tg(gap43:mGreenLantern-CAAX)* transgenic line. These fish were subjected to ONC and imaged over the course of 10 days. The lack of pigment in this transgenic line allowed us to observe *gap43* reporter signal without melanophore cell obstruction (**Figure 3A-I**). Growth cones of RGC axons were observed in the cTeO at 4 dpONC (**Figure 3H-H’ and Supplemental Video 4**). Quantification of the *gap43* reporter signal in single z-plane images from the medial TeO zone (**Figure 3A and 3B’-G’**) showed that *gap43* reporter signal increased in the cTeO at 4 dpONC, peaked at 5 dpONC, then diminished by 8-10 dpONC (**Figure 3J-K**; pre ONC vs 4 dpONC *P* value = 0.0446; pre ONC vs 5 dpONC *P* value = 0.0346). No significant difference in the *gap43* reporter signal was observed between males and females at any time-point tested (**Figure 3K**). In addition, we observed a progression of the *gap43* reporter signal in the cTeO from a rostral to caudal position over time (**Figure 3J, 3L and Supplemental Video 4**). While the downregulation of *gap43* reporter signal correlated with results from the DLR assay which showed vision recovery from 5-10 dpONC, the head tilt angle continued to improve after the complete downregulation of *gap43* reporter signal expression (**Figure S5L**). We speculate that after *gap43* reporter downregulation, additional modifications occur in the neuronal circuitry that facilitate functional recovery. There was no significant upregulation of *gap43* reporter signal in the ipsilateral optic tectum (iTeO) between pre ONC and 5 dpONC conditions (**Figure 3M-Q**), suggesting a complete decussation of the optic nerve at the optic chiasm similar to that in zebrafish (Diekmann et al., 2015). Taken together, we have generated a novel DC transgenic line that takes advantage of the *gap43* promoter to visualize *in vivo* axon regeneration. When combined with functional assays, this line has allowed us to correlate cellular phenotypes with long distance RGC axon regeneration with vision recovery.

### Regenerated axons display an altered arborization pattern post-ONC

In pro-regenerative vertebrates, studies have shown that RGC axons re-innervate the TeO after an optic nerve injury (Dow et al., 1994; Zou et al., 2013; Hu and Veldman, 2025). However, it is unclear if RGC axons re-establish connections for functional vision recovery by replicating their original axonal arborization pattern after injury. Imaging the brain of the same animal before and after injury would help answer this question but this has not been feasible using traditional adult animal models. To compare the arborization pattern of RGC axons in the TeO before injury and post-regeneration, we used sparse labeling to visualize individual cells by introducing mosaic expression of the *isl2b:TdTomato-caax* plasmid in *mitf^−/-^* fish. The zebrafish *isl2b* promoter drives transgene expression in healthy RGCs (Ben Fredj et al., 2010). We imaged the entire contralateral MB region of uninjured fish, subjected them to an ONC injury, then imaged them throughout the regeneration process (**Figure 4A**). Compared to pre-injury conditions, we observed degeneration of RGC axons at 1 dpONC as indicated by the fragmentation and reduction of the *isl2b*-driven fluorescent signal in the TeO (**Figure 4B-C and 4B’-C’**). By 5 dpONC, an extensive network of regenerating neurites was observed (**Figure 4D and 4D’**). Post-regeneration, neuronal remodeling was observed between 5-10 dpONC (**Figure 4D-E and 4D’-E’**), corresponding to the time when gradual improvements in the DLR were observed (**Figure 1D**). When detailed optical sectioning of the cTeO was performed in the same animal before and after an ONC, and the mean intensity of the signal was plotted by depth, a general similarity in the distribution of the re-innervated RGC axons was observed (**Figure 4F-H, 5A-K and Supplemental Video 5**). Interestingly, axonal remodeling continued to occur from 25-40 dpONC, a time-point after injured fish exhibit a normal DLR (**Figure 5G-K and 5J’-K’**). By comparing the patterning of RGC neurites before injury and at 40 dpONC, we observed a stark difference in the dorsal cTeO (**Figure 5L**). When imaging at a deeper medial level within the TeO, we observed the expanding arborization of regenerating RGC axons from 10-40 dpONC, a period later than that in the dorsal TeO (**Figure 5G and 5M-P**). Similar to that in the dorsal TeO, the patterning of RGC axons in the deeper medial zone of the TeO pre-injury was distinct from that at 40 dpONC (**Figure 5N**).

**Figure 5.**
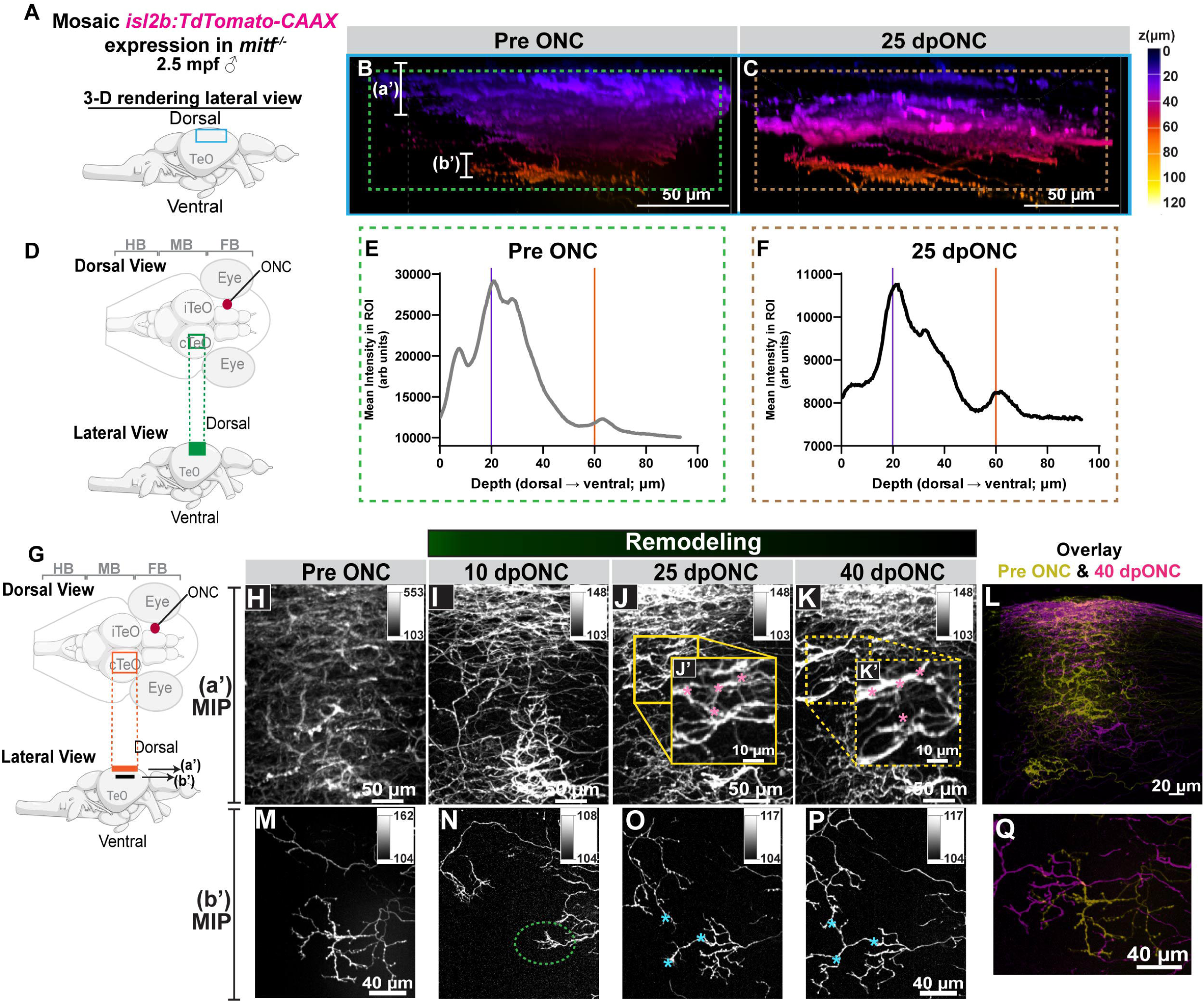
Longitudinal assessment of the re-innervation patterning of RGC axons pre-and post ONC. **(A)** Schematic representation of the lateral view of the *D. cerebrum* brain where the blue box represents the region imaged, three-dimensionally (3-D) rendered, and presented in B-C. ♂ = Male. **(B-C and H-Q)** Images were acquired using a 40X objective (N.A. = 1.15, 0.4 µm/z-step) in a male *mitf^−/-^* fish at 2.5 mpf and 1.36 cm in length with mosaic *isl2b:TdTomato-CAAX* expression. **(B-C)** Color coded 3-D rendered images of mosaic *isl2b:TdTomato-CAAX* expression at pre ONC and 25 dpONC. The total z-stack thickness was 125.6 µm and imaging z-depth is indicated in the scale. (a’) and (b’) Represent the z-depth region shown in the MIP images of (H-L) and (M-Q), respectively. **(D)** Schematic representation of the dorsal and lateral view of the ROI used for calculating the mean *isl2b:TdTomato-CAAX* signal intensity in E-F on the confocal images shown in B-C, respectively. **(E-F)** Mean *isl2b:TdTomato-CAAX* signal intensity in the ROI at different z-depths at the time-points indicated. **(G)** Schematic representation of regions imaged and shown in H-Q. The orange rectangle (a’) indicates the region shown H-K and the black rectangle (b’) indicates the region shown in M-P. **(H-K)** MIPs of the dorsal TeO region before ONC injury and at the time-points indicated. The total z-stack depth was 72.4 µm. Yellow boxes are magnified ROIs (J’ and K’). Grey scale images with LUTs shown in the upper right inset. Pink asterisks = remodeling of *isl2b:TdTomato-CAAX* expressing neurites. **(L)** Overlay of pre ONC (yellow) and 40 dpONC (magenta) images. **(M-P)** MIPs of a deeper medial TeO region before ONC injury and at the time-points indicated. The total z-stack depth was 25.6 µm. Grey scale images with LUTs shown in the upper right inset. Dotted green oval = re-forming arbor field. Blue asterisks = remodeling of *isl2b:TdTomato-CAAX* expressing neurites. **(Q)** Overlay of pre ONC (yellow) and 40 dpONC (magenta) image.

**Figure 6.**
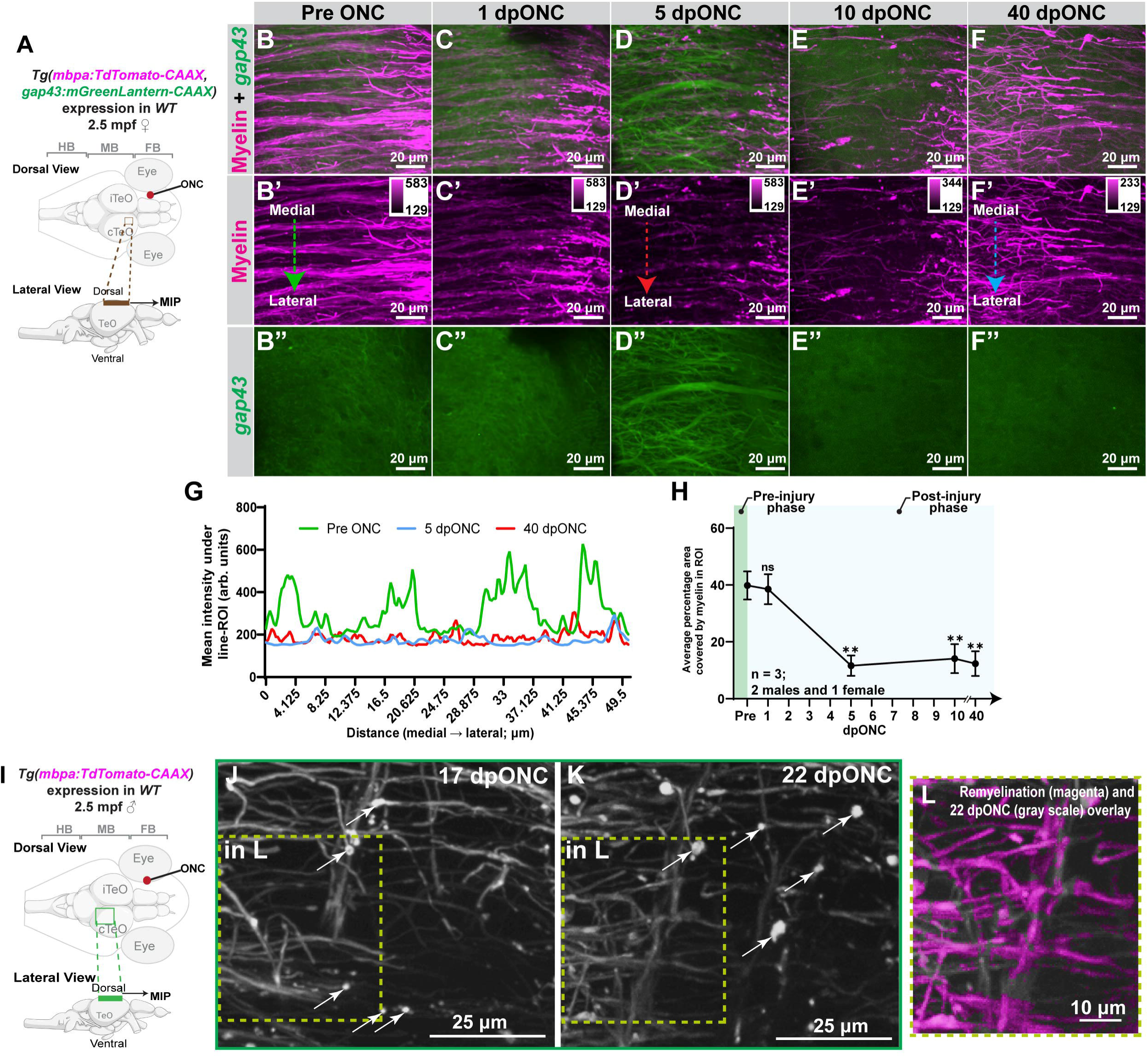
Correlating RGC axonal regeneration with oligodendrocyte membrane reporter expression in the TeO after an ONC. **(A)** Schematic representation of area imaged in B-F, B’-F’ and B’’-F’’ is indicated by the brown rectangles in the dorsal and lateral view. Images were acquired then presented as MIPs with a 40X objective (N.A. = 1.15, 0.4 µm/z-step) in a double transgenic female *Tg(mbpa:TdTomato-CAAX, gap43:mGreenLantern-CAAX)* at 2.5 mpf and 1.37 cm in length. ♀ = female **(B-F)** Merged fluorescent signals from the myelin reporter (magenta) and the gap43 reporter (green) at the time-points indicated. **(B’-F’)** Myelin reporter (magenta) only. LUTs shown in the upper right inset. (**B’’-F”)** gap43 reporter (green) only. **(G)** Mean intensity of myelin signal under a line-ROI extending from the medial to lateral cTeO as indicated in B’, D’ and F’. **(H)** Percentage area covered by *mbpa:TdTomato-CAAX* signal in a fixed ROI in the cTeO. Data represented as mean ±SEM from 3 animals (2 males and 1 female). *P* values were calculated using 1-way ANOVA (pre vs 1 dpONC *P* = 0.9987; pre ONC vs 5 dpONC *P* = 0.0055; pre ONC vs. 10 dpONC *P* = 0.0099 and pre ONC vs 40 dpONC *P* = 0.0065). Annotations as presented on graph: ns *P* ≥ 0.05, ** *P* < 0.01. **(I)** Schematic representation of the ROI imaged in a double transgenic male *Tg(mbpa:TdTomato-CAAX, gap43:mGreenLantern-CAAX)* fish, 2.5 mpf with a length of 1.27 cm, that was subjected to an ONC injury. Confocal images presented as MIPs in J-K. ♂ = male. **(J-K)** All images were acquired using a 40X objective (N.A. = 1.15, 0.4 µm/z-step). The total z-stack thickness was 33.33 µm. **(J)** Signal from the myelin reporter in the cTeO region at 17 dpONC presented in greyscale. **(K)** The same region from the same fish was imaged at 22 dpONC. (White arrows = myelin fragments). **(L)** Overlay image of remyelination (magenta; fluorescent signal subtraction of area shown in J from K) and 22 dpONC (gray scale).

### Myelinating oligodendrocyte membrane dynamics during RGC axon regeneration in the cTeO

Myelination in the CNS has been well-documented using longitudinal imaging in developing zebrafish larvae, owing to their optical transparency (Karttunen et al., 2017; Hughes and Appel, 2020; Li et al., 2024). Based on fixed time-point histological data and results from pooled tissue samples, it has been reported that adult zebrafish can restore myelin to full thickness 4 weeks after a CNS injury (Münzel et al., 2014; Zou et al., 2015). To visualize myelinating oligodendrocyte membrane dynamics and analyze the relationship between myelin and regenerating neurites in adult DC *in vivo*, we generated a double transgenic line *Tg(mbpa:TdTomato-CAAX, gap43:mGreenLantern-CAAX)* where both oligodendrocyte membranes and regenerating neurites are fluorescently labeled. Following ONC, we observed changes in the morphology of the oligodendrocyte membrane reporter signal (**Figure 6A-D, 6B’-D’ and 6B”-D”**) and its pattern (**Figure 6G**), alongside changes in regenerating neurites in the cTeO (**Figure 6B-D and 6B”-D”**). There was a significant decrease in the percentage area covered by myelin in the cTeO at 5 dpONC compared to pre-injury levels (**Figure 6H**; pre ONC vs 5 dpONC *P* value = 0.0055). At this time point, re-innervation of *gap43* positive axons in the cTeO was observed (**Figure 6D and 6D”**), with some myelin fragments in contact with regenerating neurites (**Figure S7A-C, 7B’-C’, 7B”-C” and Supplemental Video 6**). When comparing the pre-injury state (**Figure 6B and 6B”**) to later post ONC time points, the myelin pattern remained altered (**Figure 6E-F, 6E’-F’ and 6G**) and myelin levels remained reduced (**Figure 6H**; pre ONC vs 40 dpONC *P* = 0.0065). Even at 60 dpONC, myelin patterns remained distinct from that in the iTeO (**Figure S7D-F**). Although the *mbpa:TdTomato-CAAX* oligodendrocyte membrane reporter labels both demyelinating myelin fragments and newly formed myelin, we were able to detect demyelination and remyelination by comparing the myelin pattern in the same region across different time points (**Figure 6I-L**).

In conclusion, our work demonstrates that with the development of unique transgenic lines, robust longitudinal imaging techniques, and vision-based behavioral assays, adult DC can be utilized for comprehensive studies that quantify and correlate CNS axon regeneration, myelination, and functional recovery.

## Discussion

Many questions remain unanswered regarding the regeneration of axons in the CNS (Tsata and Wehner, 2021). A major hurdle has been understanding how regenerating axons re-innervate their targets in the process of reforming functional circuits. Observing this phenomenon *in vivo* and identifying contributing cellular interactions is essential. Some progress has been achieved by comparing the differential regenerative capacity of teleost fish such as zebrafish and medaka (Lai et al., 2017; Lai et al., 2019; Shimizu and Kawasaki, 2021; Chowdhury et al., 2022). However, a limitation to this approach is the inability to analyze the contribution of dynamic spatio-temporal interactions between different cell types during CNS regeneration and functional recovery. In this study, we have established a new adult animal model, *Danionella cerebrum*, for the longitudinal study of CNS regeneration using the optic nerve crush injury. We harnessed the properties of their optical transparency and small size in adulthood and performed longitudinal *in vivo* imaging of cell specific fluorescent reporters throughout the injury and regeneration process. We demonstrated that adult DC possess the ability to regenerate their RGC axons following injury, ultimately leading to functional visual recovery. Imaging revealed that axons regenerated over a long distance and exhibited evidence of axonal remodeling. Real-time *in vivo* characterization of regeneration and recovery was facilitated by generating novel DC transgenic reporter lines for axonogenesis and myelinated oligodendrocyte membranes. For longitudinal assessment of RGCs pre- and post-regeneration, we employed a sparse labeling technique using an RGC specific promoter. CRISPR/Cas9 genome editing was used to generate a pigment mutant line to enhance *in vivo* brain imaging. Additionally, two behavior-based assays were developed to assess vision recovery. Using these tools, we found that through axon remodeling events, the regenerated axon network displayed an arborization pattern distinct from its pre-injury state and that regenerating axons re-innervated appropriate targets, leading to functional vision recovery. However, levels of myelin expression and its pattern of distribution were not fully restored even well after vision recovery. The DC model provides new avenues for future studies aimed at understanding how successful CNS regeneration is orchestrated.

The time course of axonal regeneration events in the TeO following optic nerve injury was illustrated using *Tg(gap43:mGreenLantern-CAAX)* and mosaic *isl2b:TdTomato-CAAX* expression experiments. While reporter expression driven by the *gap43* promoter is transiently upregulated in regenerating axons, it is not specific to RGCs. The *isl2b* promoter, on the other hand, drives reporter expression specifically in RGCs, regardless of injury. However, literature suggests that the *isl2b* gene is downregulated following optic nerve injury in other teleost fish (Chen et al., 2022; Ali et al., 2025). Due to this property, tectal denervation and reinnervation were not quantified using the *isl2b* promoter, as it is not possible to measure and compare reporter signal intensity within the same animal at different time points to reflect the extent of RGC axon regeneration. Additionally, the transient and mosaic nature of *isl2b* reporter expression used for sparse RGC labeling prevents reliable signal quantification across animals. Given the distinct properties of the *gap43* and *isl2b* promoters, they complement each other by enabling visualization of early axon regeneration and subsequent axon reinnervation, respectively. In this study, the survival rate of RGCs in DC following ONC was not characterized. We observed that the same labeled RGCs reinnervated the optic tectum after ONC in the sparse and mosaic labeled *isl2b* reporter experiments. Due to the mosaic nature of the labeling, the regenerating axons are likely derived from the injured *isl2b* reporter expressing RGCs that were imaged prior to ONC. However, since an increase in the number of RGCs throughout life has been reported in other teleost fish (Wan et al., 2016), future studies are needed to characterize the contribution of newly formed RGCs to functional visual recovery following optic nerve injury in adult DC.

To establish a functional connection, axons must form a specific pattern to ensure proper target innervation (Gibson and Ma, 2011; Hasegawa and Kuwako, 2022). Since traditional animal models provide limited *in vivo* optical access to monitor regenerating axons in mature brains, little data is available to determine whether similar axonal re-arborization patterns are important for vision recovery. We tested the hypothesis that vision recovery relies on the post injury arborization patterns produced by regenerated axons matching their pre-injury state. We performed longitudinal imaging on animals with mosaic RGC labeling, monitored the progression of regenerating axons after injury, and compared arborization patterns between pre- and post-injury states. After ONC, we observed Wallerian degeneration followed by re-innervation of axons into the cTeO. Initially, regenerating axons formed an extensive network. Later, at a phase coinciding with vision recovery, axon remodeling occurred. Intriguingly, when comparing the axonal innervation in the same animal between the pre-injury state and the remodeling phase, we observed a similar distribution of the re-innervated neurites within the cTeO but with distinct arborization patterns. In conclusion, we demonstrated that RGC axons can regenerate and re-innervate correct targets in the TeO, leading to vision recovery.

It remains unclear, however, whether all RGCs possess the same capacity for axon regeneration. Studies using this sparse labeling approach in combination with specific promoters for RGC subpopulations will provide insight into the capacity and necessity of different RGC subtypes to regenerate in response to guidance molecules. Our work shows that the DC model, combined with our transgenic lines and imaging methods, provides a powerful platform for the *in vivo* investigation of mechanisms that regulate axonal arborization pattern formation and refinement.

The relationship between myelination and regenerative outcomes remains unclear in adult teleosts (Münzel et al., 2014; Kar:unen et al., 2017). Our data revealed that injured animals continued to exhibit reduced myelin levels in the cTeO even after vision recovery, suggesting that although functional recovery occurs relatively quickly, a return to pre-injury myelin status takes longer or may never be achieved. Although the DLR and LR assays provide information on the intactness of the animal’s vision, visual acuity remains untested. Our results suggest that vision recovery occurs before the completion of remyelination. Further investigation is needed to determine the relationship between visual acuity and remyelination. In the current study, it remains unclear whether all regenerated RGC axons are re-myelinated and whether the *mbpa:TdTomato-CAAX* oligodendrocyte membrane reporter labels compact myelin. Future investigations using a combination of different RGC axon and myelin reporters along with electron-microscopy and functional recovery assays, will provide insight into the role of myelination in different stages of axonal regeneration and functional recovery.

The DC model serves as a novel system for uncovering the fundamental mechanisms underlying CNS regeneration. It must be acknowledged that species-specific differences in CNS regeneration capabilities and anatomical features between DC and humans exists. Zebrafish have been proven to be a valuable translational research model (Phillips and Westerfield, 2014) with 71.4% of human genes having at least one zebrafish orthologue (Howe et al., 2013). A recent study suggested that 85% of zebrafish genes have a homolog in DC (Schulze et al., 2018b), highlighting the translational potential of the DC model. Currently, 86% of the DC genome is annotated (Kadobianskyi et al., 2019). Future efforts to continuously improve the DC genome sequence and its annotation will facilitate transcriptomic analyses and cross-species comparisons, enabling the identification of candidate pro-regenerative pathways for further validation in non-regenerative models.

Together, the arsenal of adult transgenic live imaging tools and visual functional recovery assays established in this study create a powerful and unique opportunity for the utilization of DC to study optic nerve regeneration. Our transgenic lines, combined with functional recovery assays, hold promise for advancing our understanding of how the nervous system is successfully rewired after injury and for gaining insight into how this process is regulated across diverse contexts — including, but not limited to, brain stab injuries, spinal cord injuries, and other injuries to sensory systems such as olfactory and auditory systems.

## Acknowledgements

Wild-type DC stocks were obtained from the laboratory of Dr. Adam Douglass (University of Utah). The *gap43* promoter containing plasmid *pW1GFG43SA* was a kind gift from Dr. Ava Udvadia (University of Wisconsin). Dr. Ross Collery (Medical College of Wisconsin) cloned the *pME-mGreenLantern* construct. Hannah G. Tebbs (Medical College of Wisconsin) performed the LR reaction and purified the *gap43:mGreenLantern-CAAX* construct. This work is funded by the National Institutes of Health Office of the Director under Award Number R21OD033664 (P.Y.L.), S10OD032136 (to Brandon Tefft - MCW engineering core), and the Advancing a Healthier Wisconsin Endowment (P.Y.L). Portions of the schematic diagrams were created using BioRender (BioRender.com).

## Author contributions

M.M. and P.Y.L. conceptualized and designed the experiments, interpreted the results, wrote and revised the manuscript with input from all authors. M.M, J.L.A, N.R.W. and J.A.P. performed the experiments, analyzed the results and wrote the Materials and Methods section. M.B.V. generated the *gap43:mGreenLantern-CAAX* plasmid and assisted in establishing the ONC technique and the DLR assay.

## Conflicting Interest

All authors declare no competing interests.

## Data and materials availability

Materials (including transgenic lines and plasmids), as well as the data supporting this study are available from the corresponding author upon request.

## Notes

***Funding:*** This work was supported by the National Institutes of Health Office of the Director under Award Number R21OD033664 (to PYL), S10OD032136 (to Brandon Tefft - Medical College of Wisconsin engineering core), and the Advancing a Healthier Wisconsin Endowment (to PYL).

### Competing Interest Statement

The authors have declared no competing interest.

### Summary of Updates

Figures, figure legends and discussion are revised.

## References

Ali A, Schriever H, Kostka D, Kuwajima T, Koenig KM, Gross J (2025) Zebrafish optic nerve injury results in systemic retinal ganglion cell dedifferentiation. bioRxiv:2025.2004.2009.646875.

Beckers A, Bergmans S, Van Dyck A, Moons L (2023) Analysis of Visual Recovery After Optic Nerve Crush in Adult Zebrafish. Methods Mol Biol 2636:437–447.

Belin S, Nawabi H, Wang C, Tang S, Latremoliere A, Warren P, Schorle H, Uncu C, Woolf CJ, He Z, Steen JA (2015) Injury-induced decline of intrinsic regenerative ability revealed by quantitative proteomics. Neuron 86:1000–1014.

Ben Fredj N, Hammond S, Otsuna H, Chien CB, Burrone J, Meyer MP (2010) Synaptic activity and activity-dependent competition regulates axon arbor maturation, growth arrest, and territory in the retinotectal projection. J Neurosci 30:10939–10951.

Bormann P, Zumsteg VM, Roth LW, Reinhard E (1998) Target contact regulates GAP-43 and alpha-tubulin mRNA levels in regenerating retinal ganglion cells. J Neurosci Res 52:405–419.

Britz R, Conway KW, Rüber L (2021) The emerging vertebrate model species for neurophysiological studies is Danionella cerebrum, new species (Teleostei: Cyprinidae). Sci Rep-Uk 11:18942–18942.

Campbell BC, Nabel EM, Murdock MH, Lao-Peregrin C, Tsoulfas P, Blackmore MG, Lee FS, Liston C, Morishita H, Petsko GA (2020) mGreenLantern: a bright monomeric fluorescent protein with rapid expression and cell filling properties for neuronal imaging. Proc Natl Acad Sci U S A 117:30710–30721.

Chato-Astrain J, García-García ÓD, Campos F, Sánchez-Porras D, Carriel V (2022) Basic Nerve Histology and Histological Analyses Following Peripheral Nerve Repair and Regeneration. In: Peripheral Nerve Tissue Engineering and Regeneration (Phillips JB, Hercher D, Hausner T, eds), pp 151–187. Cham: Springer International Publishing.

Chen S, Lathrop KL, Kuwajima T, Gross JM (2022) Retinal ganglion cell survival after severe optic nerve injury is modulated by crosstalk between Jak/Stat signaling and innate immune responses in the zebrafish retina. Development 149.

Chowdhury K, Lin S, Lai SL (2022) Comparative Study in Zebrafish and Medaka Unravels the Mechanisms of Tissue Regeneration. Frontiers in Ecology and Evolution 10.

Cook V, Groneberg AH, Hoffmann M, Kadobianskyi M, Veith J, Schulze L, Henninger J, Britz R, Judkewitz B (2024) Ultrafast sound production mechanism in one of the smallest vertebrates. Proc Natl Acad Sci U S A 121:e2314017121.

Daeschler SC, Zhang J, Gordon T, Borschel GH (2022) Optical tissue clearing enables rapid, precise and comprehensive assessment of three-dimensional morphology in experimental nerve regeneration research. Neural Regen Res 17:1348–1356.

Diekmann H, Kalbhen P, Fischer D (2015) Characterization of optic nerve regeneration using transgenic zebrafish. Front Cell Neurosci 9:118.

Dow KE, Levine RL, Solc MA, DaSilva O, Riopelle RJ (1994) Axonal transport of proteoglycans in regenerating goldfish optic nerve. Exp Neurol 126:129–137.

Dunn TW, Gebhardt C, Naumann EA, Riegler C, Ahrens MB, Engert F, Del Bene F (2016) Neural Circuits Underlying Visually Evoked Escapes in Larval Zebrafish. Neuron 89:613–628.

Gibson DA, Ma L (2011) Developmental regulation of axon branching in the vertebrate nervous system. Development 138:183–195.

Groneberg AH, Dressler LE, Kadobianskyi M, Muller J, Judkewitz B (2024) Development of sound production in Danionella cerebrum. J Exp Biol 227.

Hall TE, Martel N, Ariotti N, Xiong Z, Lo HP, Ferguson C, Rae J, Lim YW, Parton RG (2020) In vivo cell biological screening identifies an endocytic capture mechanism for T-tubule formation. Nat Commun 11:3711.

Hasegawa K, Kuwako K-i (2022) Molecular mechanisms regulating the spatial configuration of neurites. Seminars in Cell & Developmental Biology 129:103–114.

Hecker A, Anger P, Braaker PN, Schulze W, Schuster S (2020) High-resolution mapping of injury-site dependent functional recovery in a single axon in zebrafish. Communications Biology 3:307.

Howe K et al. (2013) The zebrafish reference genome sequence and its relationship to the human genome. Nature 496:498–503.

Hu M, Veldman MB (2025) Srebf2 mediates successful optic nerve axon regeneration via the mevalonate synthesis pathway. Molecular Neurodegeneration 20:28.

Hughes AN, Appel B (2020) Microglia phagocytose myelin sheaths to modify developmental myelination. Nat Neurosci 23:1055–1066.

Jung SH, Kim S, Chung AY, Kim HT, So JH, Ryu J, Park HC, Kim CH (2010) Visualization of myelination in GFP-transgenic zebrafish. Dev Dyn 239:592–597.

Kadobianskyi M, Schulze L, Schuelke M, Judkewitz B (2019) Hybrid genome assembly and annotation of Danionella translucida. Scientific Data 6:156.

Kang H (2021) Sample size determination and power analysis using the G*Power software. J Educ Eval Health Prof 18:17.

Karimi-Abdolrezaee S, Verge VM, Schreyer DJ (2002) Developmental down-regulation of GAP-43 expression and timing of target contact in rat corticospinal neurons. Exp Neurol 176:390–401.

Karttunen MJ, Czopka T, Goedhart M, Early JJ, Lyons DA (2017) Regeneration of myelin sheaths of normal length and thickness in the zebrafish CNS correlates with growth of axons in caliber. PLoS One 12:e0178058.

Kwan KM, Fujimoto E, Grabher C, Mangum BD, Hardy ME, Campbell DS, Parant JM, Yost HJ, Kanki JP, Chien CB (2007) The Tol2kit: a multisite gateway-based construction kit for Tol2 transposon transgenesis constructs. Dev Dyn 236:3088–3099.

Labun K, Montague TG, Gagnon JA, Thyme SB, Valen E (2016) CHOPCHOP v2: a web tool for the next generation of CRISPR genome engineering. Nucleic Acids Res 44:W272–276.

Labun K, Montague TG, Krause M, Torres Cleuren YN, Tjeldnes H, Valen E (2019) CHOPCHOP v3: expanding the CRISPR web toolbox beyond genome editing. Nucleic Acids Res 47:W171–W174.

Lai SL, Marín-Juez R, Stainier DYR (2019) Immune responses in cardiac repair and regeneration: a comparative point of view. Cell Mol Life Sci 76:1365–1380.

Lai SL, Marín-Juez R, Moura PL, Kuenne C, Lai JKH, Tsedeke AT, Guenther S, Looso M, Stainier DY (2017) Reciprocal analyses in zebrafish and medaka reveal that harnessing the immune response promotes cardiac regeneration. Elife 6.

Lam PY (2022) Longitudinal in vivo imaging of adult Danionella cerebrum using standard confocal microscopy. Dis Model Mech 15.

Lambert CJ, Freshner BC, Chung A, Stevenson TJ, Bowles DM, Samuel R, Gale BK, Bonkowsky JL (2018) An automated system for rapid cellular extraction from live zebrafish embryos and larvae: Development and application to genotyping. PLoS One 13:e0193180.

Lee TJ, Briggman KL (2023) Visually guided and context-dependent spatial navigation in the translucent fish Danionella cerebrum. Curr Biol 33:5467–5477 e5464.

Li J, Miramontes TG, Czopka T, Monk KR (2024) Synaptic input and Ca2+ activity in zebrafish oligodendrocyte precursor cells contribute to myelin sheath formation. Nature Neuroscience 27:219–231.

Lim JH, Stafford BK, Nguyen PL, Lien BV, Wang C, Zukor K, He Z, Huberman AD (2016) Neural activity promotes long-distance, target-specific regeneration of adult retinal axons. Nat Neurosci 19:1073–1084.

Lindemann N, Kalix L, Possiel J, Stasch R, Kusian T, Koster RW, von Trotha JW (2022) A comparative analysis of Danionella cerebrum and zebrafish (Danio rerio) larval locomotor activity in a light-dark test. Front Behav Neurosci 16:885775.

Lindsey AE, Powers MK (2007) Visual behavior of adult goldfish with regenerating retina. Vis Neurosci 24:247–255.

Lister JA, Robertson CP, Lepage T, Johnson SL, Raible DW (1999) nacre encodes a zebrafish microphthalmia-related protein that regulates neural-crest-derived pigment cell fate. Development 126:3757–3767.

Lowe DG (2004) Distinctive image features from scale-invariant keypoints. International journal of computer vision 60:91–110.

McKee A, McHenry MJ (2020) The Strategy of Predator Evasion in Response to a Visual Looming Stimulus in Zebrafish (Danio rerio). Integr Org Biol 2:obaa023.

Mensinger AF, Powers MK (2007) Visual function in regenerating teleost retina following surgical lesioning. Vis Neurosci 24:299–307.

Montague TG, Cruz JM, Gagnon JA, Church GM, Valen E (2014) CHOPCHOP: a CRISPR/Cas9 and TALEN web tool for genome editing. Nucleic Acids Res 42:W401–407.

Münzel EJ, Becker CG, Becker T, Williams A (2014) Zebrafish regenerate full thickness optic nerve myelin after demyelination, but this fails with increasing age. Acta Neuropathol Commun 2:77.

Murashova L, Dyachuk V (2025) Modeling traumatic brain and neural injuries: insights from zebrafish. Front Mol Neurosci 18:1552885.

Nevin LM, Robles E, Baier H, Scott EK (2010) Focusing on optic tectum circuitry through the lens of genetics. BMC Biol 8:126.

Oehlers SH, Cronan MR, Scott NR, Thomas MI, Okuda KS, Walton EM, Beerman RW, Crosier PS, Tobin DM (2015) Interception of host angiogenic signalling limits mycobacterial growth. Nature 517:612–615.

Park KK, Liu K, Hu Y, Smith PD, Wang C, Cai B, Xu B, Connolly L, Kramvis I, Sahin M, He Z (2008) Promoting axon regeneration in the adult CNS by modulation of the PTEN/mTOR pathway. Science 322:963–966.

Penalva-Tena A, Bedke J, Gaudin A, Barrios JP, Bertram EPL, Douglass AD Oxytocin-mediated social preference and socially reinforced reward learning in the miniature fish *Danionella cerebrum*. Current Biology.

Penalva A, Bedke J, Cook ESB, Barrios JP, Bertram EP, Douglass AD (2018) Establishment of the miniature fish species Danionella translucida as a genetically and optically tractable neuroscience model. bioRxiv.

Phillips JB, Westerfield M (2014) Zebrafish models in translational research: tipping the scales toward advancements in human health. Dis Model Mech 7:739–743.

Pinzon-Olejua A, Welte C, Chekuru A, Bosak V, Brand M, Hans S, Stuermer CA (2017) Cre-inducible site-specific recombination in zebrafish oligodendrocytes. Dev Dyn 246:41–49.

Rajan G, Lafaye J, Faini G, Carbo-Tano M, Duroure K, Tanese D, Panier T, Candelier R, Henninger J, Britz R, Judkewitz B, Gebhardt C, Emiliani V, Debregeas G, Wyart C, Del Bene F (2022) Evolutionary divergence of locomotion in two related vertebrate species. Cell Rep 38:110585.

Reinhard E, Nedivi E, Wegner J, Skene JH, Westerfield M (1994) Neural selective activation and temporal regulation of a mammalian GAP-43 promoter in zebrafish. Development 120:1767–1775.

Schaeffer J, Vilallongue N, Belin S, Nawabi H (2023) Axon guidance in regeneration of the mature central nervous system: step by step. Neural Regeneration Research 18:2665–2666.

Schindelin J, Arganda-Carreras I, Frise E, Kaynig V, Longair M, Pietzsch T, Preibisch S, Rueden C, Saalfeld S, Schmid B (2012) Fiji: an open-source platform for biological-image analysis. Nature methods 9:676–682.

Schulze L, Henninger J, Kadobianskyi M, Chaigne T, Faustino AI, Hakiy N, Albadri S, Schuelke M, Maler L, Del Bene F, Judkewitz B (2018a) Transparent Danionella translucida as a genetically tractable vertebrate brain model. Nat Methods 15:977–983.

Schulze L, Henninger J, Kadobianskyi M, Chaigne T, Faustino AI, Hakiy N, Albadri S, Schuelke M, Maler L, Del Bene F, Judkewitz B (2018b) Transparent Danionella translucida as a genetically tractable vertebrate brain model. Nature Methods 15:977–983.

Shimizu Y, Kawasaki T (2021) Differential Regenerative Capacity of the Optic Tectum of Adult Medaka and Zebrafish. Front Cell Dev Biol 9:686755.

Suthya AR, Wong CHY, Bourne JH (2024) Diving head-first into brain intravital microscopy. Front Immunol 15:1372996.

Temizer I, Donovan JC, Baier H, Semmelhack JL (2015) A Visual Pathway for Looming-Evoked Escape in Larval Zebrafish. Curr Biol 25:1823–1834.

Thuma TBT, Bogovic JA, Gunton KB, Jimenez H, Negreiros B, Pulido JS (2023) The big warp: Registration of disparate retinal imaging modalities and an example overlay of ultrawide-field photos and en-face OCTA images. PLoS One 18:e0284905.

Tsata V, Wehner D (2021) Know How to Regrow-Axon Regeneration in the Zebrafish Spinal Cord. Cells 10.

Udvadia AJ (2008) 3.6 kb genomic sequence from Takifugu capable of promoting axon growth-associated gene expression in developing and regenerating zebrafish neurons. Gene Expr Patterns 8:382–388.

Vasconcelos RO, Bolgan M, Matos AB, Van-Dunem SP, Penim J, Amorim MCP (2024) Characterization of the vocal behavior of the miniature and transparent fish model, Danionella cerebruma). J Acoust Soc Am 155:781–789.

Veith J, Chaigne T, Svanidze A, Dressler LE, Hoffmann M, Gerhardt B, Judkewitz B (2024) The mechanism for directional hearing in fish. Nature 631:118–124.

Wan Y, Almeida AD, Rulands S, Chalour N, Muresan L, Wu Y, Simons BD, He J, Harris WA (2016) The ciliary marginal zone of the zebrafish retina: clonal and time-lapse analysis of a continuously growing tissue. Development 143:1099–1107.

Watanabe S, Takabayashi A, Takagi S, von Baumgarten R, Wetzig J (1989) Dorsal light response and changes of its responses under varying acceleration conditions. Adv Space Res 9:231–240.

Williams PR, Benowitz LI, Goldberg JL, He Z (2020) Axon Regeneration in the Mammalian Optic Nerve. Annu Rev Vis Sci 6:195–213.

Zada D, Schulze L, Yu JH, Tarabishi P, Napoli JL, Milan J, Lovett-Barron M (2024) Development of neural circuits for social motion perception in schooling fish. Curr Biol 34:3380–3391 e3385.

Zou S-q, Yin W, Huang Y-b, Tian C, Ge S-c, Hu B (2015) Chapter 2 - Functional Regeneration and Remyelination in the Zebrafish Optic Nerve. In: Neural Regeneration (So K-F, Xu X-M, eds), pp 21–41. Oxford: Academic Press.

Zou S, Tian C, Ge S, Hu B (2013) Neurogenesis of retinal ganglion cells is not essential to visual functional recovery after optic nerve injury in adult zebrafish. PLoS One 8:e57280.

