## Supplemental figures for "*In vivo* imaging of central nervous system regeneration using *Danionella cerebrum*"

*Supplemental figures and legends:*

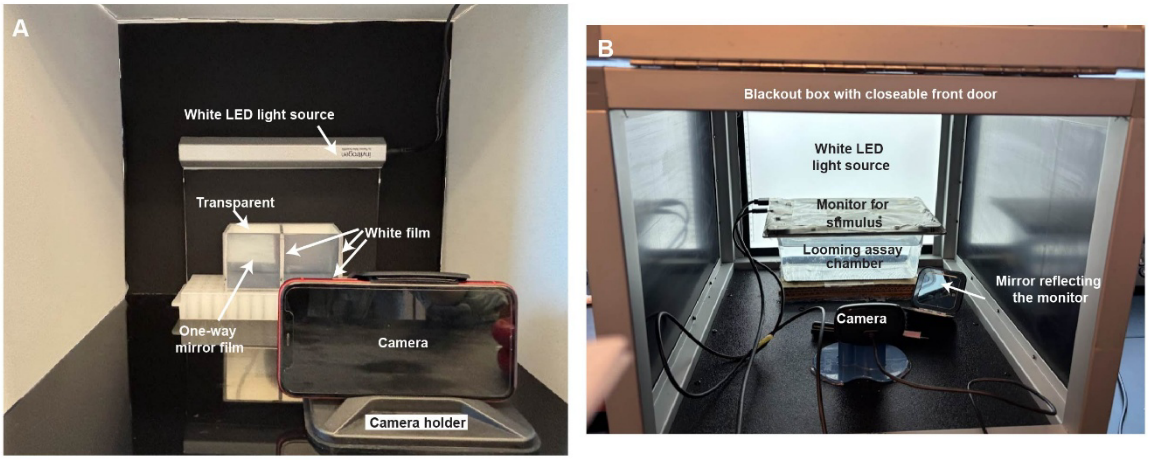


**Figure S1. Apparatus for the Dorsal Light Response (DLR) and Looming Response (LR) Assays (related to Figure 1). (A)** The DLR assay uses a 2-chamber (dimensions of 1 chamber = 6 cm x 6 cm x 6 cm) container which was covered by an opaque white film on 3 sides. The back of the DLR chamber facing the white LED light source was transparent. The front of the DLR chamber was covered with a 1-way mirror film. A camera was positioned on a stand in front of the DLR chamber. **(B)** The LR assay was performed in a blackout box with a closeable front door, inside which the LR chamber was positioned with a white LED source in the back. A monitor, used to present the LR stimulus to the fish, was positioned on the top of the LR chamber. A camera was positioned on a stand in front of the LR chamber. A mirror reflecting the presence of the LR stimulus to the camera was positioned on the right of the LR chamber.

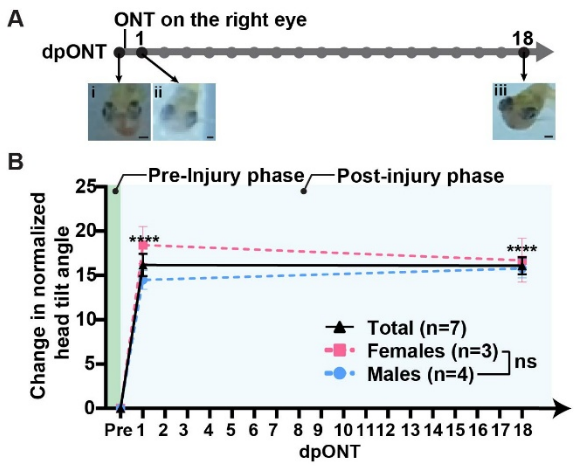


**Figure S2. Assaying vision recovery after ONT using the DLR assay (related to Figure 1).** (A) Schematic of the timeline used for the ONT experiment. The black dots indicate on what days the DLR assay was performed. (i-iii) Representative images of the same fish at (i) pre ONT, (ii) 1-day post ONT (dpONT) and (iii) 18 dpONT. Scale bar = 200 μm **(B)** The change in head tilt angle, normalized to the pre ONT angle, was plotted for the days tested. All data represented in mean ±SEM. Black solid line represents all animals (n = 7), pink dotted line represents the females (n = 3), blue dotted line represents the males (n = 4). *P* value of all animals (total) at each time-point as compared to pre ONT was calculated using Mixed Effects Repeated Measures 2-way ANOVA. *P* values from left to right: $P < 0.0001$ and $P < 0.0001$. For males vs females, at 1 dpONT $P = 0.3415$; and at 18 dpONT $P = 0.9415$. Annotations as presented on graph: **** $P < 0.0001$.

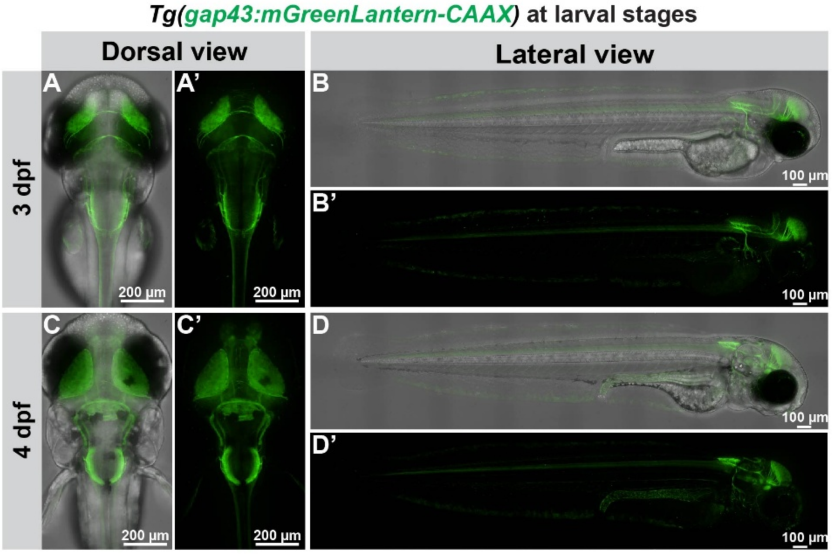


**Figure S3.** ***gap43*** **reporter expression in** ***Tg(gap43:mGreenLantern-CAAX)*** **larvae (related to Figure 2).**

**(A-D)** *gap43* reporter expression (green channel) merged with the corresponding bright field image at the time-point indicated. **(A'-D')** *gap43* reporter expression only (green channel) at the time-point indicated. (A-A') (C-C') Maximum intensity projection (MIP) images of the dorsal view. The total z-stack was 168.30 µm in (A-A') and 226.80 µm in (C-C'). (B-B') (D-D') MIP images of the lateral view. The total z-stack was 199.80 µm. All confocal images were acquired using a 20X objective (N.A. = 0.75, 0.9 µm/step).

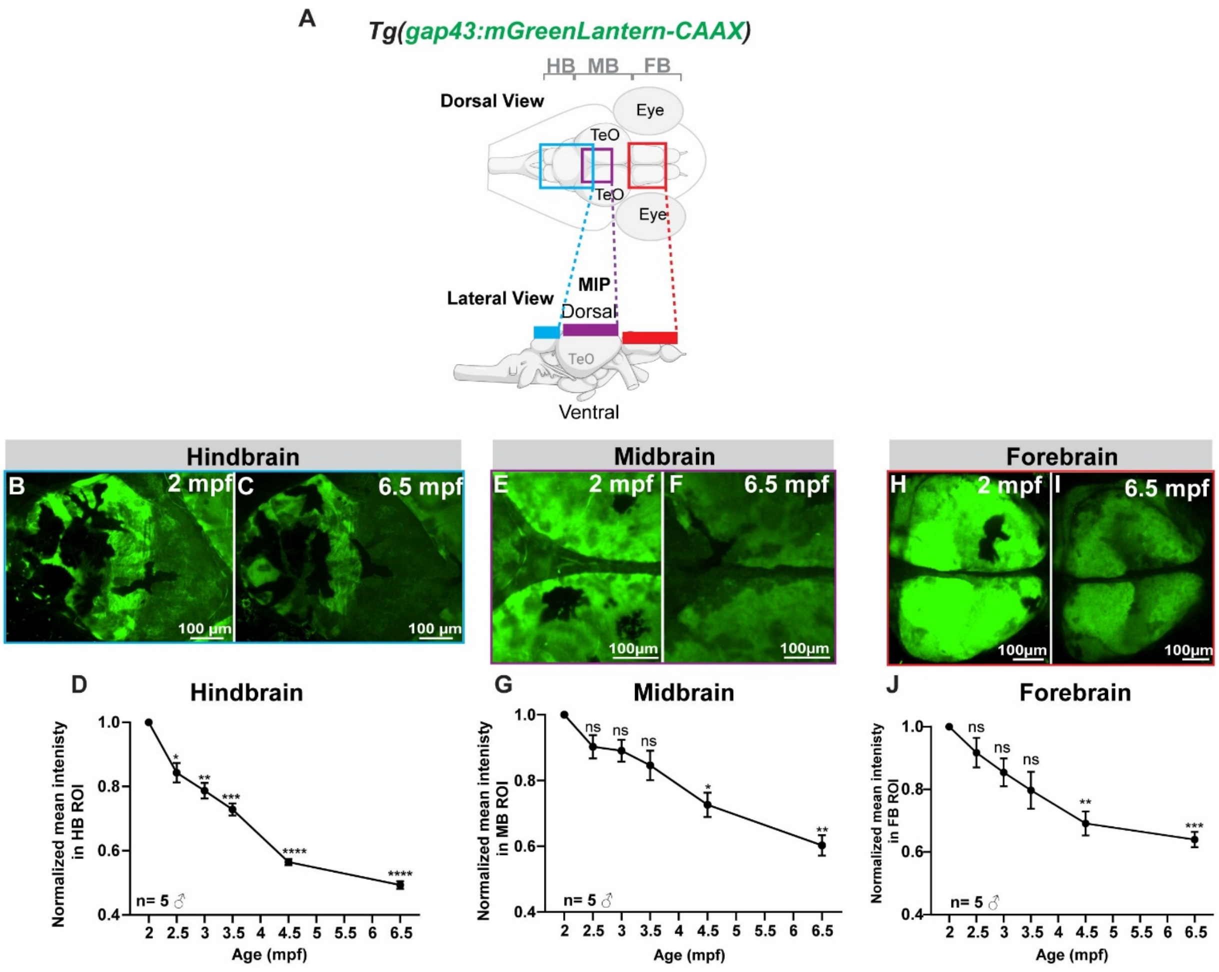


**Figure S4. Quantifying levels of *gap43* reporter expression in adult DC.**

**(A)** Illustration showing the regions imaged in the hindbrain (B-C), midbrain (E-F), and forebrain (H-I). The colored boxes indicated the approximate areas covered in each region. All confocal images were acquired with a 20X objective (N.A. = 0.75, 1.8 µm/z-step) and presented as maximum intensity projections (MIPs). ♂ = Male. **(B-C)** Dorsal view MIPs of the HB at 2 and 6.5 mpf. The total z-stack was 110 µm thick. **(D)** Quantification of mean signal intensity from gap43-driven expression (normalized to levels at 2 mpf) in adult transgenic animals. All data represented as mean ±SEM (n = 5). *P* values of each time-point were calculated against the 2 mpf time-point using Repeated Measures 1-way ANOVA (*P* values from left to right: $P = 0.0357$, $P = 0.0055$, $P =$

0.0008, $P < 0.0001$, and $P < 0.0001$). Annotations as presented on graph: ns $P \geq 0.05$, * $P < 0.05$, ** $P < 0.01$, *** $P < 0.001$, **** $P < 0.0001$. **(E-F)** Dorsal view MIPs of the MB at 2 and 6.5 mpf. The total z-stack was 95.4 µm thick. **(G)** Quantification of mean signal intensity from gap43-driven expression (normalized to levels at 2 mpf) in adult transgenic animals. All data represented as mean ±SEM (n = 5). *P* values of each time-point were calculated against the 2 mpf time-point using Repeated Measures 1-way ANOVA (*P* values from left to right: $P = 0.2412$, $P = 0.1490$, $P = 0.1348$, $P = 0.0104$, and $P = 0.0012$). Annotations as presented on graph: ns $P \geq 0.05$, * $P < 0.05$, ** $P < 0.01$. **(H-I)** Dorsal view MIPs of the FB at 2 and 6.5 mpf. The total z-stack was 228.6 µm thick. **(J)** Quantification of mean signal intensity from gap43-driven expression (normalized to levels at 2 mpf) in adult transgenic animals. All data represented as mean ±SEM (n=5). *P* values of each time-point was calculated against the 2 mpf time-point using Repeated Measures one-way ANOVA (*P* values from left to right: $P = 0.5718$, $P = 0.1548$, $P = 0.1351$, $P = 0.0076$, and $P = 0.0007$). Annotations as presented on graph: ns $P \geq 0.05$, ** $P < 0.01$, *** $P < 0.001$.

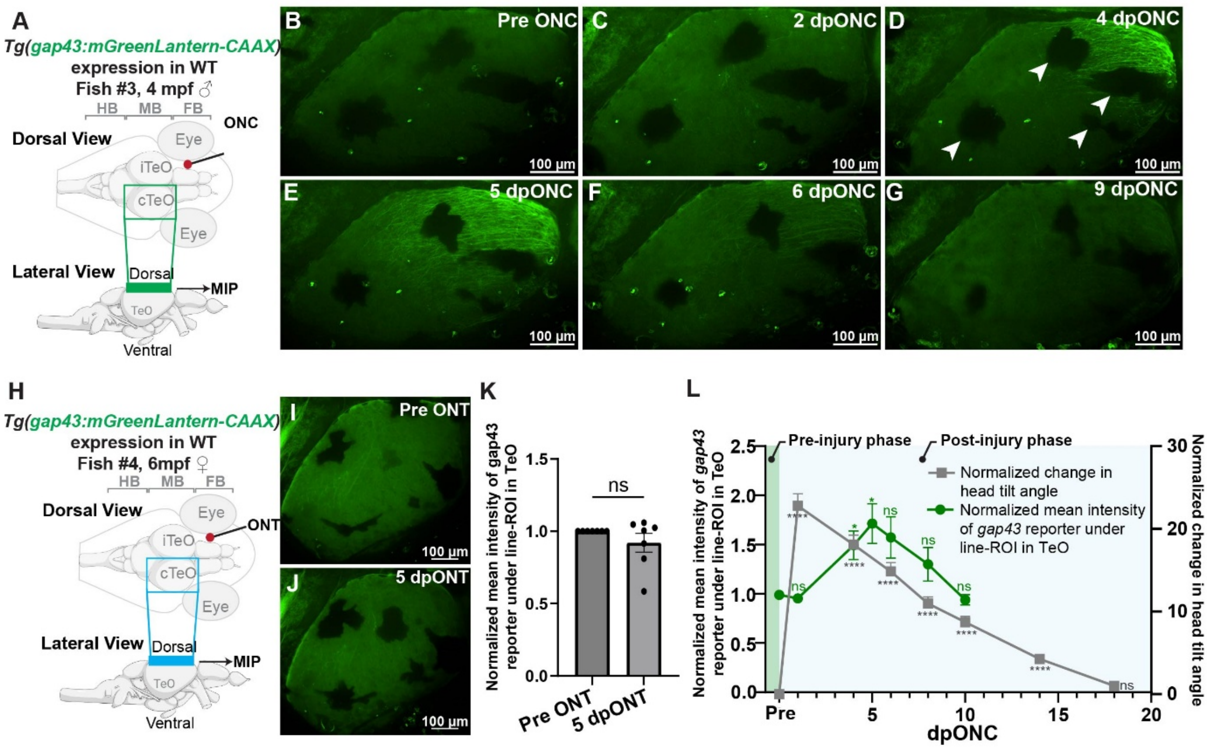


**Figure S5. Characterizing *gap43* reporter expression after optic nerve injuries (related to Figure 3).**

**(A)** Schematic representation of brain region imaged in B-G before and after an ONC injury. ♂ = Male. **(B-G)** Confocal maximum intensity projection (MIP) images at the time-points indicated were acquired using a 10× objective (N.A. = 0.45, 2.5 µm/z-step) in a *Tg(gap43:mGreenLantern-CAAX)* male at 4 mpf and 1.35 cm in length. White arrowheads in (D) indicate pigment cells. **(H)** Schematic representation of brain region imaged in J-K before and 5-days post optic nerve transection (dpONT) injury. ♀ = female **(I-J)** MIP images acquired at the time-point indicated using a 10× objective (N.A. = 0.45, 2.5 µm/z-step) in a *Tg(gap43:mGreenLantern-CAAX)* female at 6 mpf and 1.50 cm in length. **(K)** Mean intensity of *gap43* expression in the TeO before and 5 dpONT. Values were normalized to pre ONT readings and expressed as arbitrary units. All data represented as mean ±SEM (n = 6). *P* value was calculated using a 2 Tailed Paired t-test where *P* = 0.2689. Annotations as presented on graph: ns *P* ≥ 0.05. **(L)** Correlation between *gap43* expression and head tilt angles measured using the DLR assay before and after an ONC injury. Values were normalized to pre ONC readings and expressed as arbitrary units (gap43 reporter expression) or angle (head tilt). (n = 16, the same group of fish were used for *gap43* and DLR analysis). All data represented as mean ±SEM. *P* values were calculated using Mixed Effects Repeated Measures 2-way ANOVA. *P* values represented as compared to pre ONC. For the gray line: *P* values from left to right: $P < 0.0001$, $P < 0.0001$, $P < 0.0001$, $P < 0.0001$, $P < 0.0001$, $P < 0.0001$, and $P = 0.5676$. For the green line: *P* values from left to right: $P = 0.9728$, $P = 0.0446$, $P = 0.0346$, $P = 0.1417$, $P = 0.5474$, and $P = 0.9576$. Annotations as presented on graph: ns $P \geq 0.05$, * $P < 0.05$, **** $P < 0.0001$.

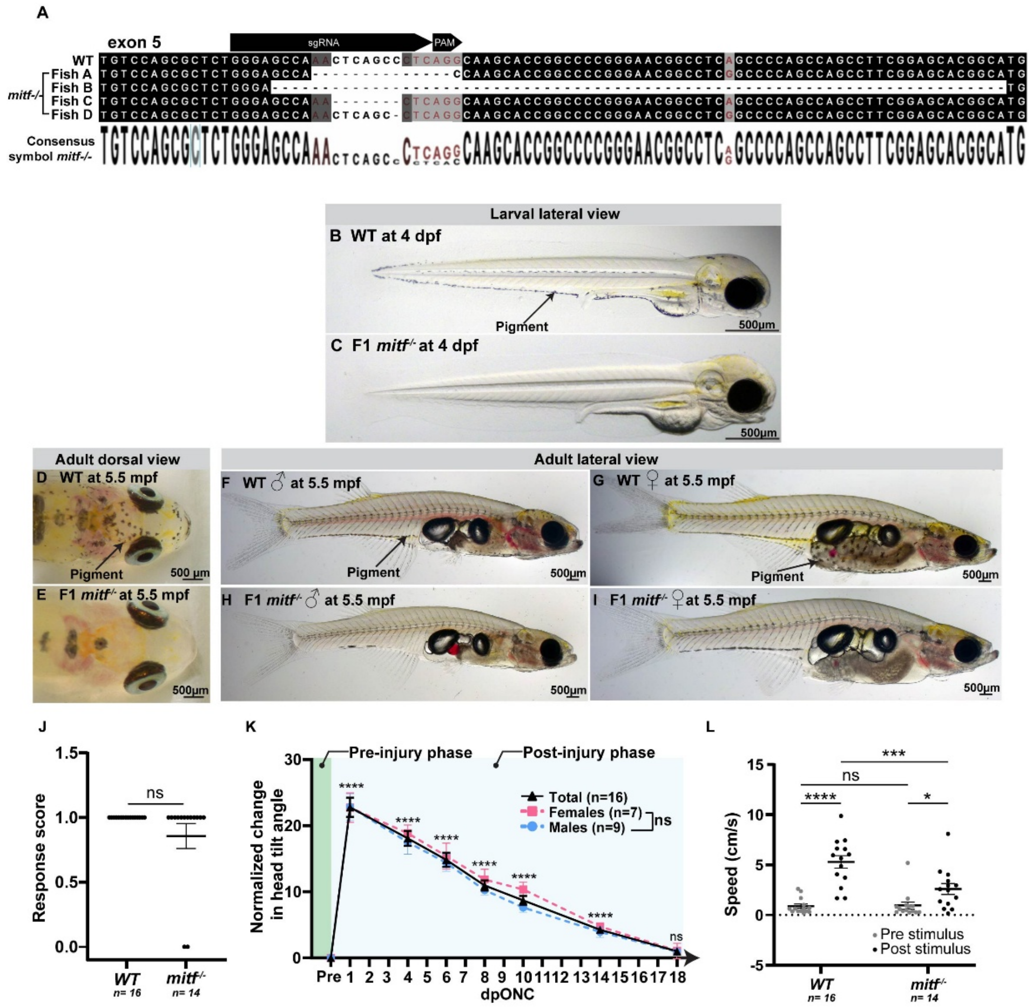


**Figure S6. Generating a DC *mitf* $^{-/-}$ pigment deficient line and evaluating vision in adult mutant fish.**

**(A)** Targeted region within exon 5 of the *mitf* gene. Sequence alignment comparing the *WT mitf* sequence to indels of isolated (founder) *mitf*$^{-/-}$fish. The sequence of the targeting gRNA is indicated. **(B)** Lateral view of larval *WT* DC at 4 dpf. Black arrow indicates the pigment spots on the ventral side of the larva. **(C)** Lateral view of larval *mitf*$^{-/-}$ DC at 4 dpf. **(D)** Dorsal view of an adult *WT* DC at 5.5 mpf. Black arrow indicates a pigment spot. **(E)** Dorsal view of an adult *mitf*$^{-/-}$ DC at 5.5 mpf. (**F-G)** Lateral view of an adult *WT* DC male (F) and female (G) at 5.5 mpf. Black arrow indicates pigment. **(H-I)** Lateral view of an adult *mitf*$^{-/-}$ DC male (H) and female (I) at 5.5

mpf. ♀ = female and ♂ = Male **(J)** Results of the LR assay comparing *WT* to *mitf*$^{-/-}$ fish. All data represented in mean ±SEM (n = 16 *WT* and n = 14 *mitf*$^{-/-}$). *P* value was calculated using 2 Tailed Unpaired t-test where $P = 0.1259$. Annotations as presented on graph: ns $P \geq 0.05$. **(K)** DLR assay performed on adult *mitf*$^{-/-}$ fish. Head tilt angle was measured before injury and over 18 dpONC. Post injury values were normalized to pre ONC levels. All data represented in mean ±SEM. Black solid line represents all animals (n = 16), pink dotted line represents the females (n=7), blue dotted line represents the males (n = 9). *P* values were calculated using Mixed Effects Repeated Measures 2-way ANOVA. *P* values of all animals (total) at each time-points compared to pre ONC from left to right: $P < 0.0001$, $P < 0.0001$, $P < 0.0001$, $P < 0.0001$, $P < 0.0001$, $P < 0.0001$, and $P = 0.5676$. For females vs males, at 1 dpONC, $P = 0.9997$; at 4 dpONC, $P = 0.8123$; at 6 dpONC, $P = 0.9025$; at 8 dpONC, P = 0.6330; at 10 dpONC $P = 0.1571$; at 14 dpONC $P = 0.6844$; and 18 dpONC $P = 0.9912$. Annotations as presented on graph: ns $P \geq 0.05$, **** $P < 0.0001$. **(L)** Swim speed (cm/s) was averaged over 1s after the LR stimulus in *WT* and *mitf*$^{-/-}$ fish. All data represented in mean ±SEM (n=16 *WT* and n=14 *mitf*$^{-/-}$). *P* value was calculated using Repeated Measures 2-Way ANOVA. *WT* pre vs post stimulus, $P < 0.0001$; *mitf*$^{-/-}$ pre vs post stimulus, $P = 0.0103$; pre stimulus *WT* vs *mitf*$^{-/-}$, $P = 0.9050$; and post stimulus *WT* vs *mitf*$^{-/-}$, $P = 0.0001$. Annotations as presented on graph: Annotations as presented on graph: ns $P \geq 0.05$, * $P < 0.05$, *** $P < 0.001$, **** $P < 0.0001$.

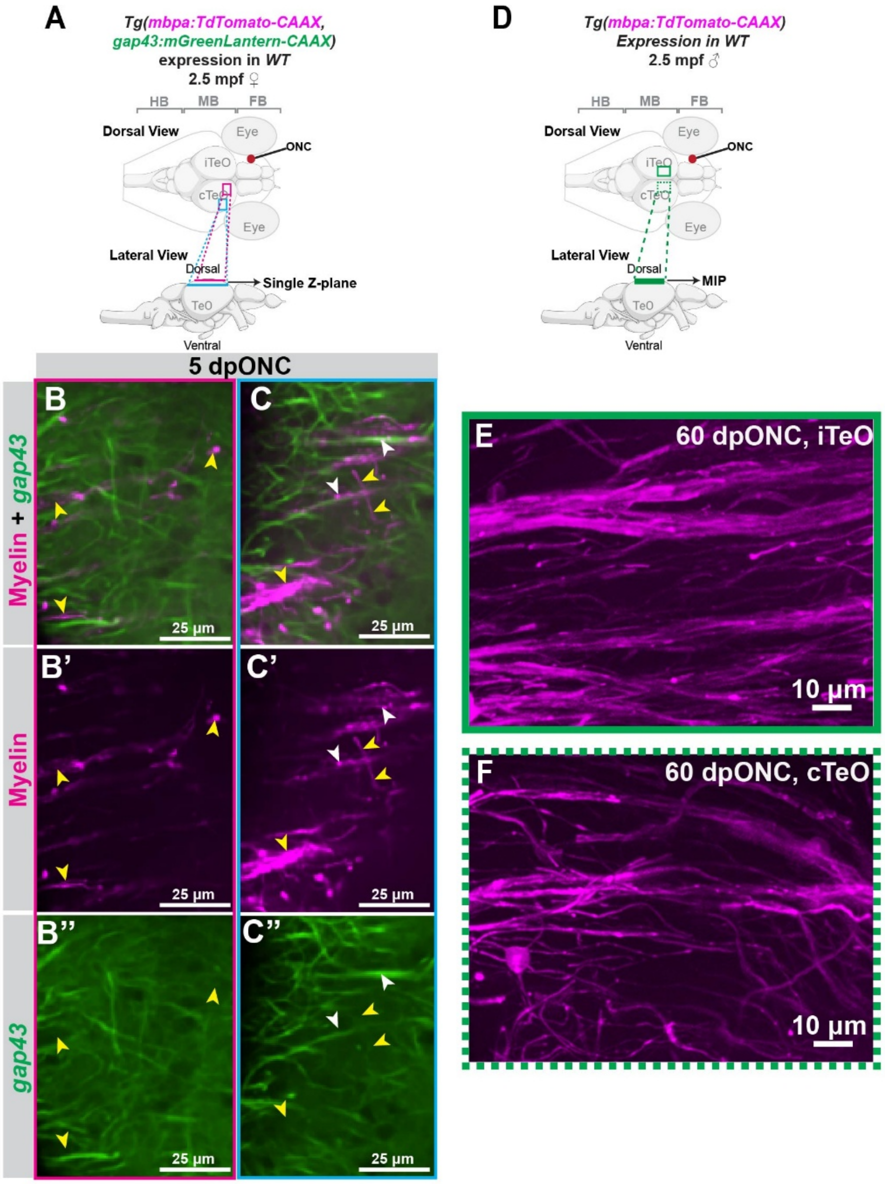


**Figure S7. Myelination in the optic tectum after an ONC injury.**

(A) Schematic representation of the regions imaged in B-C, B'-C', and B''-C''. All images are single z-planes in the area and depths indicated. ♀ = female, and ♂ = male. **(B-C)** Merged fluorescent signals from the myelin reporter (magenta) and the *gap43* reporter (green). **(B'-C')** Myelin reporter (magenta) only. **(B''-C'')** *gap43* reporter (green) only. White arrowheads indicate regions of contact between both myelin and *gap43* reporter expression. Yellow arrow heads indicate regions where only myelin reporter expression is seen. **(D)** Schematic illustration of the brain regions imaged and shown in E and F. Green boxes indicate the approximate areas imaged. Images presented as maximum intensity projections (MIPs). **(E-F)** Images were acquired at 60 dpONC with a 40X objective (N.A. = 1.15, 0.4 μm/z-step) in a male *Tg(mbpa:TdTomato-CAAX)* at 4.5 mpf 1.37 cm in length. The total z-stack used for MIP was 56.8 μm.

***Supplemental video legends:***

**Supplemental video 1: DLR of the same fish before and after an ONC injury (related to Figure 1).** Example videos showing the DLR taken before injury and at 1 and 18 dpONC in an adult female DC *WT* fish (4 mpf, 1.50 cm in length). Timecode on the top right-hand side corner shows zero based time from when the video starts (HH:MM:SS:FF).

**Supplemental video 2: LR Assay of the same fish before and after an ONC injury (related to Figure 1).** Example videos showing the LR taken before injury and at 1 and 12 dpONC in an adult female DC *WT* fish (6 mpf, 1.50 cm in length). The LR stimulus video (shown to the fish in real time) is displayed in the top left-hand corner. LR stimulus comes on at 10 seconds. Timecode on the top right-hand side corner shows zero based time from when the video starts (HH:MM:SS:FF).

**Supplemental video 3: Dorsal to ventral confocal z-series images of an adult *Tg(gap43:mGreenLantern-CAAX)* DC brain (related to Figure 2).** An adult *WT* male DC fish (2 mpf, 1.31 cm in length) was imaged using a 10X objective for a 455 μm z-depth (2.5 μm/z-step). Z-position is shown on the top right-hand corner of the image.

**Supplemental video 4: A time-lapse video of regenerating axons 4 dpONC, in the contralateral optic tectum (related to Figure 3).** Confocal z-stack images were acquired at 15 min intervals during live imaging of an adult DC *mitf*$^{-/-}$ *Tg(gap43:mGreenLantern-CAAX)* male (6-months post fertilization, 1.28 cm in length) at 4 dpONC with a 40X objective for an 80 μm Z-depth (0.4 μm/z-step). Timecode on the top right-hand side corner shows zero based time from when the video starts (HH:MM:SS). White boxes mark different growth cones over time.

**Supplemental video 5: 3-D rendered movie of the contralateral midbrain with mosaic expression of isl2b:TdTomato-CAAX (related to Figure 4).** Confocal z-stack images were acquired from an adult DC *mitf*$^{-/-}$ male (2.5-months post fertilization, 1.36 cm in length), sparsely expressing the transgene isl2b:TdTomato-CAAX. In vivo images were acquired before injury and 25 dpONC (supplemental to Fig. 5) using a 40x objective for a 125.6 µm Z-depth (0.4 µm/z-step). Images are displayed in color with depth encoding.

**Supplemental video 6: 3-D rendered movie of regenerating RGC neurites (green) and myelin (magenta) in the contralateral optic tectum region of the midbrain (related to Figure 5).** Confocal z-stack images were acquired from an adult *WT Tg(gap43:mGreenLantern-CAAX, mbpa:TdTomato-CAAX)* DC female (2.5 mpf, 1.37 cm in length) *in vivo* at 5 dpONC (supplemental to Fig. S7 A-C) using a 40X objective for a 60.40 µm Z-depth (0.4 µm/z-step).
